# The cryo-electron microscopy structure of the human CDK-activating kinase

**DOI:** 10.1101/2020.05.13.094755

**Authors:** Basil J. Greber, Juan M. Perez-Bertoldi, Kif Lim, Anthony T. Iavarone, Daniel B. Toso, Eva Nogales

## Abstract

The human CDK-activating kinase (CAK), a complex composed of cyclin dependent kinase (CDK) 7, cyclin H, and MAT1, is a critical regulator of transcription initiation and the cell cycle. It acts by phosphorylating the C-terminal heptapeptide repeat domain of the RNA polymerase II subunit Rpb1, which is an important regulatory event in transcription initiation by Pol II, and it phosphorylates the regulatory T-loop of CDKs that control cell-cycle progression. Here, we have determined the three-dimensional structure of the catalytic module of human CAK, revealing the structural basis of its assembly and providing insight into CDK7 activation in this context. The unique third component of the complex, MAT1, substantially extends the interaction interface between CDK7 and cyclin H, explaining its role as a CAK assembly factor, and it forms interactions with the CDK7 T-loop, which may contribute to enhancing CAK activity. We have also determined the structure of the CAK in complex with the covalently bound inhibitor THZ1 in order to provide insight into the binding of inhibitors at the CDK7 active site and aid in the rational design of therapeutic compounds.

**Significance:** Control of gene expression and the cell cycle is critical for appropriate cell growth and timely cell division. Failure of the mechanisms regulating these processes can result in proliferative diseases. A molecular complex termed the CDK activating kinase (CAK) impinges on both of these regulatory networks in human cells and is thus a possible drug target for treatment of cancer. Here, we use cryo-electron microscopy to describe the detailed molecular structure of the human CAK, revealing its architecture and the interactions between its regulatory elements. Additionally, we have obtained the structure of the CAK in complex with a small-molecule inhibitor.

## Introduction

Transcription initiation by eukaryotic RNA polymerase II (Pol II) depends on the formation of a pre-initiation complex (PIC) on promoter DNA, which includes general transcription factors, Pol II, and Mediator (Sainsbury et al., 2015). Like many other cellular processes, transcription and the assembly of the PIC are regulated by phosphorylation events. During PIC assembly, mediator interacts with the C-terminal heptapeptide repeat domain of Rpb1, the largest subunit of Pol II (Myers et al., 1998), also referred to as the Pol II-CTD. Phosphorylation of this region by the cyclin-dependent kinase (CDK) 7 subunit of transcription initiation factor IIH (TFIIH) disrupts its interactions with Mediator, allowing initiating Pol II to break free from the PIC and clear the promoter (Søgaard & Svejstrup, 2007). Pol II-CTD phosphorylation and de-phosphorylation are thus key events that regulate transcription of eukaryotic protein-coding genes and have been implicated in the formation of phase-separated transcriptional sub-compartments (Harlen & Churchman, 2017; Jeronimo et al., 2013). This process is not only important for normal transcription, but also contributes to aberrantly elevated transcription levels in cancer cells. Consequently, transcription-related CDKs have been identified as promising drug targets (Galbraith et al., 2019), and inhibition of CDK7 by compounds such as THZ1 or SY-1365 has been shown to selectively kill cancer cells that require elevated levels of transcription (Hu et al., 2019; Kwiatkowski et al., 2014).

In addition to its conserved role in transcription, human CDK7 phosphorylates CDKs that control the cell cycle, a function that is not conserved in yeast. The activity of cell cycle CDKs is generally controlled by association with partner cyclins (Morgan, 1997). Cyclin binding provides the bulk of CDK stimulation (Connell-Crowley et al., 1993) by inducing conformational changes in the CDK that lead to extension of a regulatory loop near the active site, known as the T-loop, and to the proper arrangement of the active site for phosphoryl-transfer (Jeffrey et al., 1995). However, in addition to cyclin binding, full activation of cell cycle CDKs requires phosphorylation of the T-loop (Morgan, 1997; Russo et al., 1996). In animal cells, these activating phosphorylations are carried out by CDK7 (Fisher & Morgan, 1994; Larochelle et al., 1998), itself a cyclin-dependent kinase whose activity depends on cyclin H (Fisher & Morgan, 1994).

In human and other metazoan cells, regulation of transcription initiation by phosphorylation of the Pol II-CTD and phosphorylation of cell cycle CDKs thus depend on a single kinase, CDK7, which functions in a complex termed the CDK-activating kinase (CAK) (Fisher & Morgan, 1994). In addition to CDK7 and cyclin H, human CAK comprises a third subunit, MAT1 (Devault et al., 1995; Fisher et al., 1995; Tassan et al., 1995; Yee et al., 1995). CDK7 and cyclin H form a canonical CDK-cyclin pair, while MAT1 serves as a CAK assembly factor (Devault et al., 1995; Fisher et al., 1995; Yee et al., 1995), increases CAK activity (Busso et al., 2000; Fisher et al., 1995), and tethers the CAK to the core of TFIIH when this ten-subunit complex carries out its function in transcription initiation (Busso et al., 2000; Serizawa et al., 1995; Shiekhattar et al., 1995).

While X-ray crystal structures of CDK7 and of cyclin H have been determined (Andersen et al., 1997; Kim et al., 1996; Lolli et al., 2004), structural information on the entire three-subunit CAK, which is critical for a mechanistic understanding of the role of MAT1 in CAK assembly and the regulation of its activity, has remained elusive. Here, we present cryo-electron microscopy (cryo-EM) structures of human CAK bound to either ATPγS or the inhibitor THZ1, thereby revealing the architecture of the human CAK and the structural basis for its inhibition by covalent inhibitors. Our structure of the CAK also completes the high-resolution structural analysis of human TFIIH and shows how MAT1 interlinks CDK7, XPB, and XPD, the three major functional centres in TFIIH.

## Results and Discussion

### Structure determination of the human CAK

To determine the structure of the human CAK, we recombinantly co-expressed CDK7, cyclin H, and MAT1 in insect cells, purified the resulting complex by affinity and size-exclusion chromatography, and collected cryo-EM data of frozen-hydrated CAK bound to the nucleotide analogue ATPγS (Fig. S1) using a 200 kV instrument (Herzik et al., 2019). Three-dimensional (3D) reconstruction resulted in a structure at 2.9-Å resolution (Fig. 1A, B, Fig. S2A-C), which allowed unambiguous docking of the high-resolution crystal structures of CDK7 and cyclin H (Andersen et al., 1997; Kim et al., 1996; Lolli et al., 2004). We rebuilt the docked models according to the cryo-EM map, which showed clear density for amino acid side chains and the bound nucleotide (Fig. 1C, S2D, E). After assignment and rebuilding of the CDK7-cyclin H dimer, the remaining density allowed building of residues 244-308 of MAT1, consistent with biochemical data showing that a C-terminal fragment of MAT1 (residues 229-309) is sufficient for binding to CDK7-cyclin H and to enhance kinase activity of CDK7 (Busso et al., 2000). The reconstructed volume thus corresponds to a mass of approximately 84 kDa, which is among the smallest asymmetric particles for which better than 3-Å resolution have been obtained using cryo-EM.

**Figure 1.**
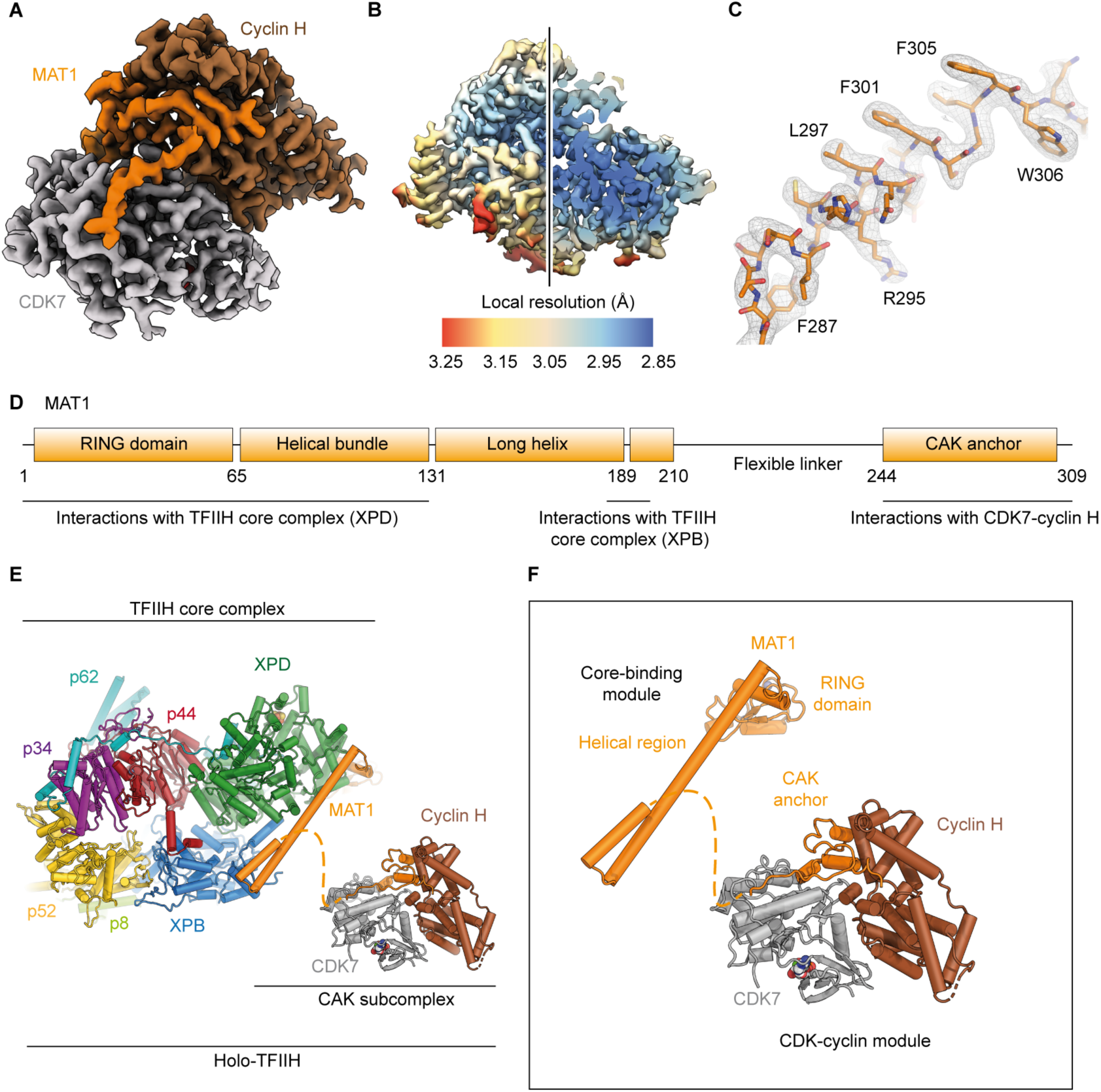
Structure determination of CAK and structural models of holo-TFIIH and the full CAK complex. (**A**) Cryo-EM map of human CAK at 2.9-Å resolution. Subunits are colored and indicated. (**B**) Local resolution estimation. (**C**) Assignment of MAT1 according to features of the cryo-EM density. (**D**) Schematic of domain organization and interactions of MAT1 with partners within TFIIH. (**E**) Depiction of holo-TFIIH, composed of the core complex (PDB ID 6NMI) and the CAK subcomplex. (**F**) The complete CAK subcomplex consists of a core-binding module (visualized with the TFIIH core complex) and the CDK-cyclin module (reported in this work).

### Complete structural description of human TFIIH

The CAK subcomplex is an important functional unit of TFIIH and critical for its activity as a general transcription factor. Previous cryo-EM studies of human and yeast TFIIH have revealed the structure of the TFIIH core complex and residues 1-210 of MAT1 (Greber et al., 2017; Greber et al., 2019; He et al., 2016; Kokic et al., 2019; Schilbach et al., 2017), but were lacking the remainder of the CAK module due to disorder (Greber et al., 2017; Greber et al., 2019) or low local resolution (He et al., 2016; Schilbach et al., 2017). Our structure of human CAK completes the structure of human TFIIH (Fig. 1D-F). The combined structures show how MAT1 simultaneously interacts with both the TFIIH core complex and the CDK7-cyclin H module. Distinct and independently folding structural modules, the RING domain and helical regions near N-terminus, and the CAK anchor near the C-terminus of MAT1, engage with the TFIIH core complex and the CDK-cyclin module, respectively. These regions are connected by a flexible tether of approximately 35 residues that are not resolved in any of the structures reported (Fig. 1D-F). The flexible nature of the tether and the lack of observed interactions of the CDK7-cyclin H module with other regions of MAT1 in the full-length CAK complex (Fig. 1A) explain why the CDK7-cyclin H module and parts of MAT1 bound to it were not resolved in previous structures of free TFIIH (Greber et al., 2017; Greber et al., 2019). This structural flexibility may support the ability of the CAK module to occupy distinct positions in the Pol II-PIC depending on the presence of mediator (He et al., 2016; Schilbach et al., 2017; Yan et al., 2019). Our findings also underscore the role of MAT1 as a key regulator of TFIIH that physically links the DNA translocase XPB, the helicase XPD, and the kinase CDK7 (Fig. 1E), the three ATP-consuming, functional centres of the complex. MAT1 forms interactions with regulatory elements on both CDK7 (see below) and XPD (Greber et al., 2019; Kokic et al., 2019; Peissert et al., 2020), highlighting its central regulatory role.

### Role of MAT1 in CAK architecture and stability

Overall, the arrangement of CDK7 and cyclin H resembles that of other CDK-cyclin complexes (reviewed in Wood & Endicott, 2018). Interactions are mostly confined to the N-terminal domains of both CDK7 and cyclin H (Fig. 2A, S3A), burying approx. 1030 Å^2^ of accessible surface area. Previous analysis revealed that both the extent of the interaction surface and the geometry of CDK-cyclin complexes show marked variability (Echalier et al., 2010). We find that the C-terminal lobe of CDK7 and the C-terminal cyclin fold of cyclin H are rotated away from each other compared to the structure of CDK2-cyclin A (Jeffrey et al., 1995), resulting in a geometry that is intermediate between that of CDK2-cyclin A and that described for the transcriptional CDK9-cyclin T complex (P-TEFb) (Baumli et al., 2008) (Fig. 2B, S3B, C). In the CAK, the C-terminal CAK-anchor region of MAT1 (residues 255-309) fills the space vacated by this rotation (Fig. 2A, B). Most of this region of MAT1 is sandwiched between the C-terminal lobe of CDK7 and the C-terminal cyclin fold of cyclin H, thereby bridging the two subunits (Fig. 2C, D). Indeed, MAT1 buries more surface area with CDK7 and cyclin H (> 1100 Å^2^ each) than CDK7 and cyclin H bury with each other. This rationalizes biochemical data that established MAT1 as a CAK assembly factor that enhances the stability of the CDK7-cyclin H interaction (Fisher et al., 1995; Yee et al., 1995), but requires revision of a previously proposed architecture of the CAK based on low-resolution cryo-EM data, which placed the MAT1 components such that the space between CDK7 and cyclin H was left unoccupied (Yan et al., 2019).

**Figure 2.**
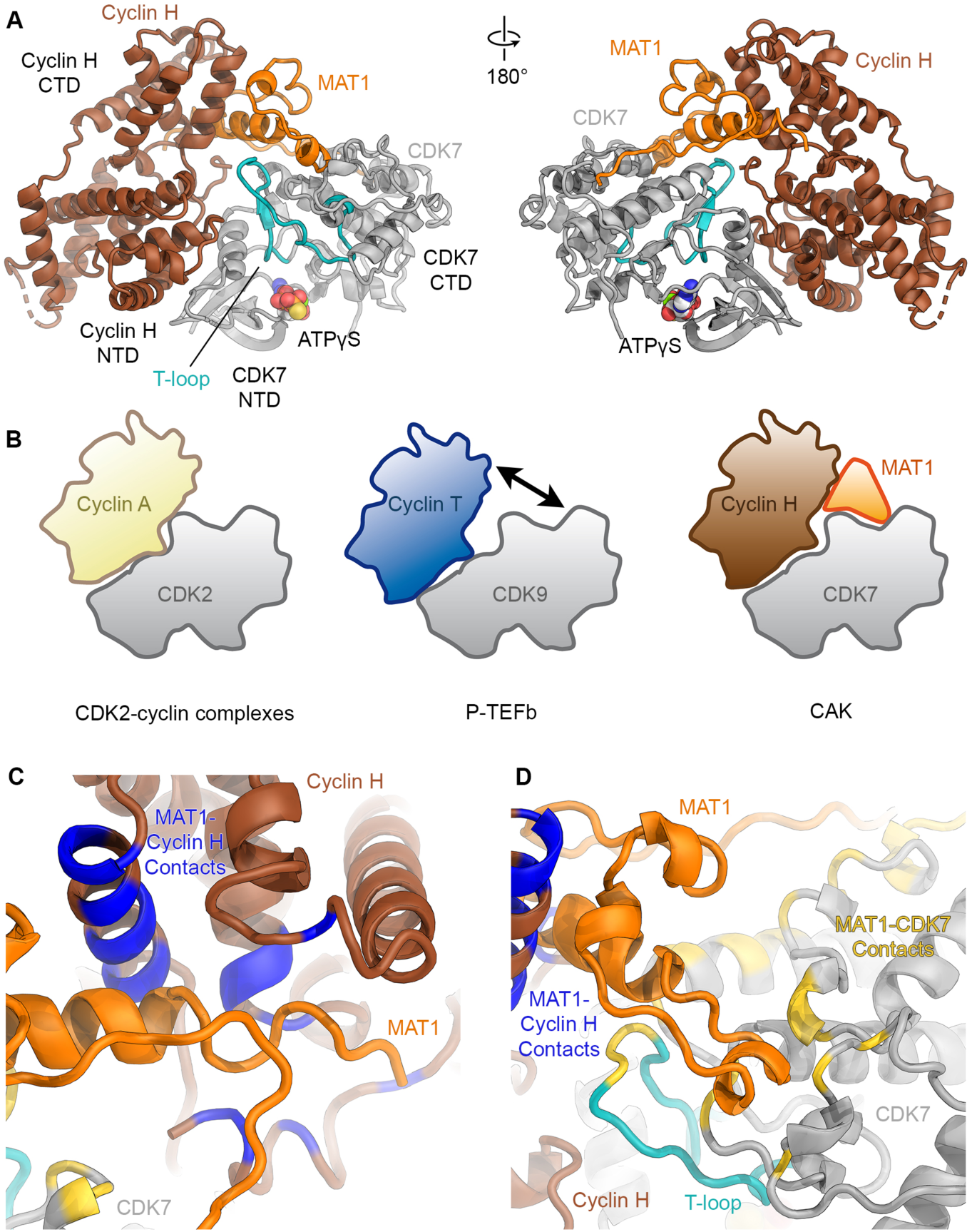
Structure of the human CAK and role of MAT1 in its architecture. (**A**) Atomic model of CAK shown in “front” and “back” views (CDK7 grey with the T-loop shown in teal, cyclin H brown, MAT1 orange). (**B**) Schematics of the architecture of CDK2-containing complexes, transcriptional CDK complexes, and CAK. (**C, D**) Interactions of MAT1 with cyclin H and CDK7. Contact sites (distance < 4 Å) on cyclin H are coloured blue, contact sites on CDK7 are coloured yellow, the T-loop is shown in cyan.

The most prominent and well-ordered part of the MAT1 density is its C-terminal helix (Fig. 1C, residues 288-300), which spans the distance between the C-terminal lobe of CDK7 and the C-terminal cyclin fold of cyclin H and interacts with both proteins. Comparison of the CAK structure to that of CDK2 with cyclins A, B, and E (Brown et al., 2007; Honda et al., 2005; Jeffrey et al., 1995; Russo et al., 1996) shows that the C-terminal helix of MAT1 occupies a similar position as an extended helix at the N-terminus of the cyclin folds in cyclin A, B, and E that contributes to the enlarged interaction surface in CDK2-cyclin complexes (Fig. S3D). Both cyclin H and cyclin T within P-TEFb (Baumli et al., 2008), which is more closely related to cyclin H than are the cell cycle cyclins (Ma et al., 2013), lack this extended helix at the N-terminus of the cyclin folds, leaving room for other interactions at this site. However, it is unclear whether any associated factors bind at this site in P-TEFb or other transcriptional CDKs, and thus the co-option of this interaction site by an associated protein may be unique to CAK.

The yeast equivalent of the human CDK7-Cyclin H-MAT1 complex is called TFIIK and is formed by the proteins Kin28, Ccnh, and Tfb3 (Feaver et al., 1994; Keogh et al., 2002). Analysis of our structure shows that the MAT1 residues that form contacts to Cyclin H and CDK7 are mostly conserved in Tfb3 (Fig. S4, S5), suggesting that the molecular contacts between MAT1/Tfb3 and the CDK-cyclin pair are likely equivalent, and confirming a conserved architecture between CAK and TFIIK, and likely across all eukaryotes.

In summary, our data establish that MAT1 forms extensive, conserved interfaces with both CDK7 and cyclin H, explaining its role as a CAK assembly factor.

### Conformation and interactions of the CDK7 T-loop in the human CAK

In our structure, which has clear density for ATPγS in the CDK7 active site (Fig. S2D), the regulatory T-loop of CDK7 is found in an extended, active conformation, pointing away from the nucleotide-binding site and towards cyclin H (Fig. 3A, B, Fig. S6). Accordingly, our purified CAK was able to phosphorylate a synthetic (YSPTSPS)_3_KKK peptide, mimicking the RPB1 CTD, on Ser5 of the YSPTSPS heptapeptide repeat (data not shown). Notably, the tip of the CDK7 T-loop closely approaches the C-terminal helix of MAT1 (Fig. 2D), suggesting that interactions with MAT1 might promote the extended conformation of the CDK7 T-loop within the CAK complex (Fig. 3A, C), likely contributing to the reported CAK activation by MAT1 (Busso et al., 2000).

**Figure 3.**
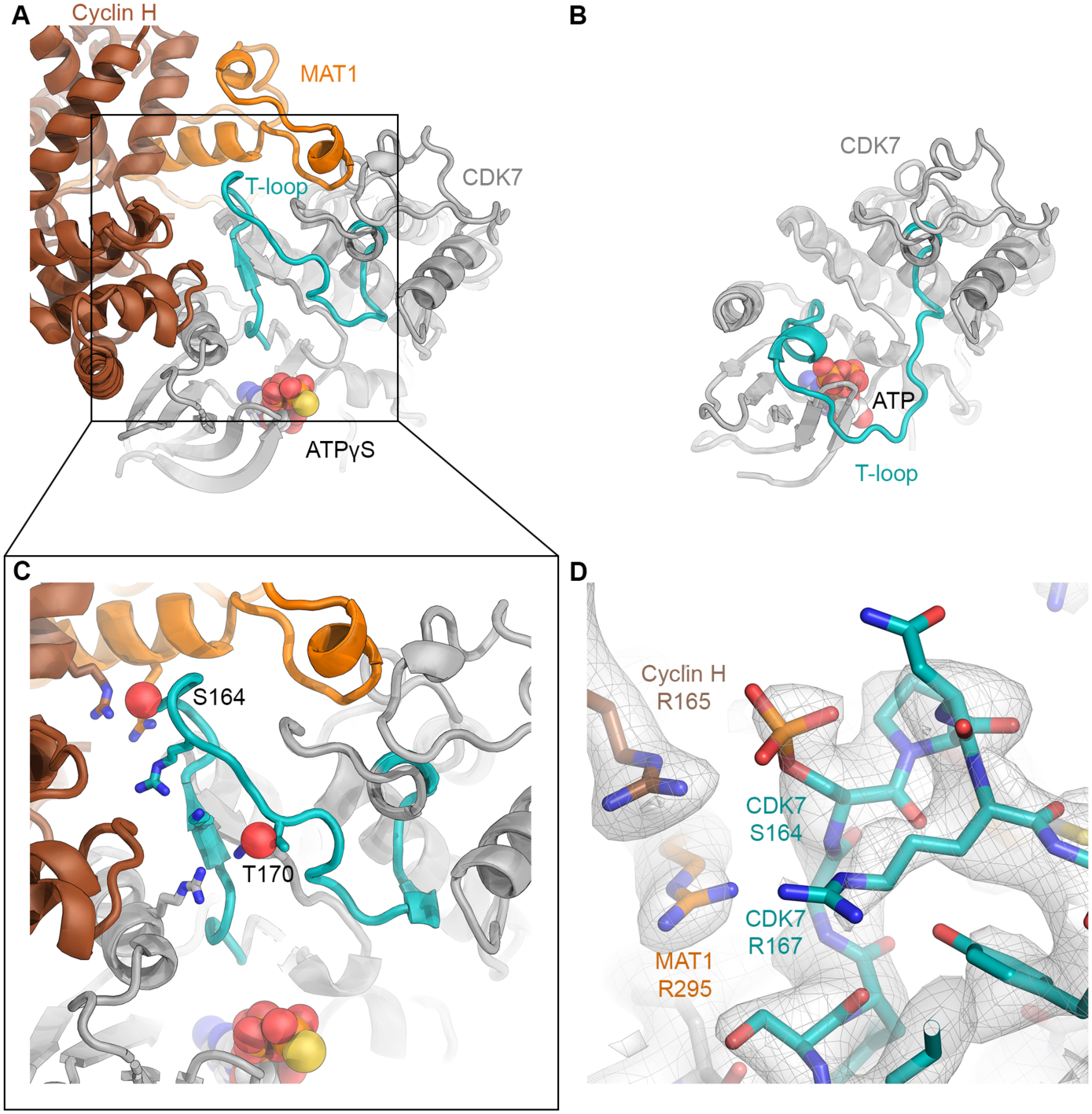
Extended T-loop of CDK7. (**A**) The T-loop in the CAK-ATPγS complex is extended and points towards MAT1 and cyclin H. (**B**) In the crystal structure of free, inactive CDK7, the T-loop is folded across the active site (PDB ID 1UA2; shown in the same orientation as CAK in panel A) (Lolli et al., 2004). (**C**) Positively charged residues near the CDK7 phosphorylation sites (S164, T170; Oγ shown as red spheres) are shown as sticks. (**D**) Structure of the CAK near S164; cryo-EM density shown in grey (combined surface and mesh) suggests that S164 is partially phosphorylated in our preparation of CAK. Positively charged residues surrounding the phosphate group are labeled.

Like its CDK substrates, CDK7 itself is the target of regulatory phosphorylation. Specifically, its T-loop harbors residues S164 and T170 (Fig. 3C), both of which are phosphorylation sites (Fisher et al., 1995; Labbé et al., 1994; Martinez et al., 1997). In our cryo-EM maps, S164 and to a lesser degree T170 showed enlarged densities that may arise from the presence of sub-stoichiometric levels of the phosphorylated species (Fig. 3D). These map features are consistent with mass spectrometric analysis indicating the presence of some phosphorylated molecules in our CAK samples (Fig. S7). The presence of both phosphorylated and unphosphorylated CDK7 in otherwise homogeneously stable CAK is in agreement with previous reports of MAT1-mediated assembly of CAK showing that full, active complex can exist in the absence of T-loop phosphorylation (Fisher et al., 1995).

Interestingly, S164 within the CDK7 T-loop is found in proximity of three arginine side chains (Fig. 3C, D), one each from all three CAK subunits (CDK7 R167, MAT1 R295, cyclin H R165). This patch of arginines is reminiscent of a positively charged pocket that accommodates the phosphate group of phosphorylated T160 in CDK2, an arrangement that was first visualized in the structure of phosphorylated CDK2 in complex with cyclin A (Russo et al., 1996), and which is also found in CDK7 (CDK7 residues R61, R136, and K160; Fig. 3C). The human CAK thus harbors two such positively charged pockets, the first of them formed by CDK7 alone and conserved across CDKs, the second specific to the CAK and formed by all three subunits. Phosphate binding in the latter pocket might influence complex assembly and/or the conformation of the T-loop. Indeed, we found that a phosphate group modelled into our density closely approaches cyclin H R165, suggesting formation of an intermolecular salt bridge, and the phosphate may form additional interactions with CDK7 R167 and N166 (Fig. 3D). These potential additional intermolecular interactions following phosphorylation of S164 might promote CAK stability or an extended conformation of the T-loop, both of which would likely contribute to regulation of CAK activity towards certain substrates. This idea is consistent with previous findings showing that T-loop phosphorylation can stabilize CDK7-cyclin H association in the absence of MAT1, and that an S164A mutant shows lower kinase activity towards CDK2 (Fisher et al., 1995). Additionally, an S164A T170A double mutant shows reduced CAK assembly in *Drosophila* (Larochelle et al., 2001) and an equivalent mutant in *Xenopus laevis* CDK7 (S170A) shows impaired activation in response to cyclin H binding (Martinez et al., 1997). Our structural results can rationalise these biochemical data. However, our structure does not provide a clear mechanism for the inhibition of CTD-kinase activity by S164 phosphorylation (Akoulitchev & Reinberg, 1998), and it cannot easily explain why T170 phosphorylation was found to contribute more strongly to CAK assembly than S164 phosphorylation (Fisher et al., 1995; Martinez et al., 1997), given that T170 is far removed from any structural elements of cyclin H or MAT1 (Fig. 3C). It is possible that T170 phosphorylation has an indirect effect by affecting CDK7 conformation, or that it stabilizes an extended T-loop conformation that enables formation of interactions involving the tip of the T-loop.

Taken together, our data suggest that T-loop contacts with MAT1 and formation of intermolecular salt bridges by phosphorylated S164 are specific features of the CAK that may contribute to an extended state of the T-loop and thus promote a CDK7 conformation that is active towards its CDK targets.

### Structural mechanism of irreversible inhibition of CDK7 by THZ1

Due to its activity in transcription and cell cycle regulation, the human CAK plays an important role in regulating cell growth and division. Consequently, inhibition of the CAK has been found to be a promising strategy for treatment of proliferative diseases (Galbraith et al., 2019; Kwiatkowski et al., 2014). To obtain insight into the covalent binding of the cysteine-reactive anti-cancer compound THZ1 (Kwiatkowski et al., 2014) to CDK7 within the CAK, we have determined the 3.4-Å-resolution structure of THZ1 bound to a CDK7-cyclin H-MAT1Δ219 complex (CAK-MAT1Δ219), in which the N-terminal region of MAT1, including its RING domain, has been removed (Fig. 4A, S1A, S8). Consistent with mass spectrometric analysis that shows essentially complete occupancy of our complex with THZ1 (Fig. S7), we observe well-defined density for the small-molecule inhibitor in the active site of CDK7 (Fig. 4B). THZ1 occupies the mostly hydrophobic binding pocket that otherwise accommodates the base and ribose of bound ATP (Fig. 4C-E). Less defined density extends towards the comparably poorly ordered C-terminal region of CDK7, which is only visible until residue C312. In inhibitor-bound CDK7, the nucleotide binding pocket is slightly contracted at the site where the phosphate groups would be located in the nucleotide-bound state (Fig. 4F), likely because there are no corresponding groups in the inhibitor occupying this position.

**Figure 4.**
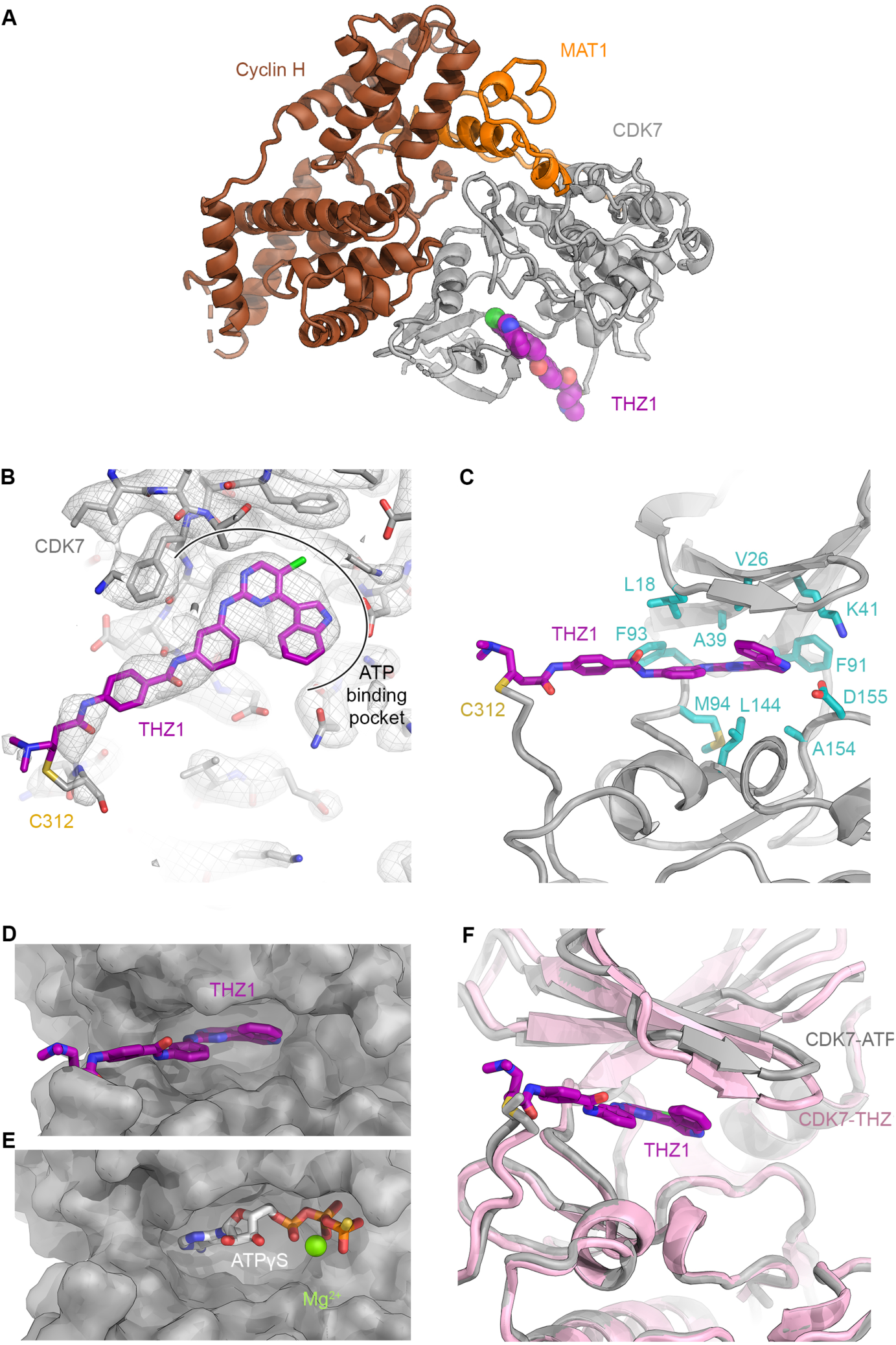
The CAK-THZ1 complex. (**A**) Structure of the CAK-THZ1 complex. THZ1 is shown in purple. (**B**) THZ1 shown in the cryo-EM density contoured at 4.5σ. THZ1 is bound in the ATP-binding pocket of CDK7 and stretches to the covalently modified cysteine C312. (**C**) Molecular environment of THZ1 in the nucleotide binding pocket of CDK7, dominated by hydrophobic residues. (**D**) THZ1 (purple) in the nucleotide binding cavity of CDK7. (**E**) ATPγS in the nucleotide binding cavity of CDK7. The aromatic head group of THZ1 and the base and ribose of the nucleotide occupy the same site. (**F**) Comparison of the conformation of CDK7 in the nucleotide-bound (grey) and THZ1-bound (pink) states.

Our observations are in overall agreement with the proposed mode of binding of THZ1 based on computational modelling, and with the observed reactivity of the THZ1 acrylamide group towards C312 in CDK7 (Kwiatkowski et al., 2014). Our density is best defined for the section of THZ1 that is buried inside CDK7 and suggests that the THZ1 indole ring is rotated such that it points towards the open side of the nucleotide binding pocket, rather than more deeply into it (Fig. 4B, C). This aspect of the conformation of THZ1 more closely resembles a recent computational model for the binding of SY-1365 to CDK7 (Hu et al., 2019), as compared to the originally proposed model for binding of THZ1 (Kwiatkowski et al., 2014). In summary, while supporting the previous computational model for the interaction of THZ1 with CDK7 at the ATP binding pocket, and in agreement with the formation of an irreversible adduct outside the active site, our structure now refines our understanding of the conformation of the inhibitor inside the CDK7 nucleotide binding pocket.

## Conclusion

The cryo-EM structure of the 84-kDa CDK-cyclin module of the human CAK reveals the molecular architecture of this trimeric CDK assembly, how the unique third subunit, MAT1, contributes to the stability of the complex, and how molecular interactions within the CAK promote the extended conformation of the T-loop of CDK7. We also determined the structure of the small-molecule inhibitor near the active site of CAK. Our study shows that cryo-EM is suitable for structure determination of a small, asymmetric complex at better than 3-Å resolution, opening the way for structure-guided design and improvement of therapeutic compounds that inhibit the activity of the human CAK and other important targets of similar size.

## Methods

### Cloning and expression of CAK and CAK-MAT1Δ219

Codon-optimized genes for MAT1, cyclin H, and CDK7 (sequences were synthesized by GenScript) were amplified by polymerase chain reaction (PCR) using primers with suitable overhangs and cloned into 438-series vectors harboring expression cassettes with baculoviral promoters and terminators (Gradia et al., 2017) using the in-fusion variant (Irwin et al., 2012) of SLIC cloning (Li & Elledge, 2012). The expression cassettes encoding His_6_-tagged MAT1, STREP-tagged CDK7, and tag-free cyclin H (Fig. S1A) were then combined into expression constructs containing MAT1-Cyclin H-CDK7 using PmeI (MssI) and SwaI (SmiI) restriction digests (FastDigest enzymes, ThermoFisher Scientific) and subsequent in-fusion reactions. The construct for expression of MAT1Δ219-cyclin H-CDK7 (CAK-MAT1Δ219) was assembled using the same approach but using cloning primers that remove the N-terminal 219 residues of MAT1 and allow creation of an N-terminal MBP-His_6_-fusion construct. A construct harboring a Pol II-CTD peptide mimic fused to the CDK7 N-terminus was constructed by linearization of the plasmid encoding CDK7 by PCR, followed by insertion of a gene block (synthesized by IDT DNA) encoding the peptide sequence and a linker by an in-fusion reaction. The peptide could not be visualized in any of the subsequent cryo-EM maps (see below).

The vectors encoding the expression constructs were transformed into EMBacY cells (Bieniossek et al., 2008) for preparation of bacmids by standard methods. Bacmid DNA was prepared by isopropanol precipitation and transfected into Sf9 (*Spodoptera frugiperda*) insect cells using FuGENE HD transfection reagent (Promega). After two rounds of virus amplification in Sf9 cells, 1-L cultures of High5 (*Trichoplusia ni*) insect cells were infected with 10 mL supernatant from the previous virus amplification cultures. The cells were harvested by centrifugation after 72 hours of incubation and frozen in liquid N_2_ for later use.

### Purification of CAK

CAK complexes were purified using Nickel affinity, Strep-Tactin affinity, and size exclusion chromatography. Cells were resuspended in lysis buffer (250 mM KCl, 40 mM HEPES-KOH pH 7.9, 5 mM MgCl_2_, 5 mM β-mercaptoethanol, 10 mM imidazole and 10% (v/v) glycerol supplemented with protease inhibitors and DNaseI) and homogenized by sonication. The lysate was cleared using a JA-20 rotor (18,000 rpm for 30 min at 4°C) and incubated with Ni-NTA superflow resin (Qiagen) for 30 min. The beads were washed in purification buffer (250 mM KCl, 25 mM HEPES-KOH pH 7.9, 5 mM MgCl_2_, 5 mM β-mercaptoethanol, and 10% (v/v) glycerol) supplemented with 25 mM imidazole and eluted in purification buffer with 300 mM imidazole. The eluate was incubated with Strep-Tactin superflow plus resin (Qiagen) for 30 min, washed with 10 bed volumes of purification buffer, and eluted with purification buffer supplemented with 10 mM desthiobiotin (Sigma-Aldrich). The eluted fractions were concentrated by ultrafiltration using 30-kDa molecular weight cut-off Centricons centrifugal filter units (EMD Millipore) and applied onto a Superdex 200 10/300 GL column (GE Healthcare) for size exclusion chromatography (Fig. S1B). Purified fractions (Fig. S1C) were pooled, concentrated to 2 mg/mL, and frozen in liquid N_2_ for storage at -80°C.

### Purification of CAK-MAT1Δ219 and THZ1 complex formation

For studies involving THZ1 binding to CAK, the CAK-MAT1Δ219 construct was used because THZ1 induced precipitation of full-length CAK at various stages during the preparation of the complex, possibly because the zinc-coordinating cysteines in the MAT1 RING domain can engage in unwanted reactions with THZ1. CAK-MAT1Δ219 was purified as described above, with modifications to the protocol to enable binding of THZ1. Specifically, the MBP-His_6_ tag on MAT1 and the Strep-tag on CDK7 were cleaved using TEV protease (2 h at room temperature). The resulting protein was incubated with nickel beads to remove the cleaved MBP-tag and TEV protease and subsequently subjected to gel filtration to remove β-mercaptoethanol. This step was added to avoid side-reactions of the sulfhydryl-reactive acrylamide group in THZ1 with the thiol on β-mercaptoethanol. After formation of the CAK-MAT1Δ219-THZ1 complex by incubation of 0.2 mg/mL CAK-MAT1Δ219 (approx. 2 μM) with 5 μM THZ1 (EMD Millipore) for 1.5 h at room temperature, the complex was concentrated and subjected to a second gel filtration step, again without β-mercaptoethanol, to remove excess THZ1 and residual DMSO, in which THZ1 had been solubilized. The complex was then concentrated, flash frozen and stored at -80°C.

### Cryo-EM specimen preparation

Cryo-EM specimens were prepared on UltrAuFoil R1.2/1.3 holey gold grids (Quantifoil microtools) that were plasma cleaned using a Tergeo plasma cleaner (PIE Scientific) for 15-45 sec. 2 mg/mL CAK was diluted 10x in 200 mM KCl, 2 mM MgCl_2_, 20 mM HEPES-KOH pH 7.9, 5 mM β-mercaptoethanol and incubated for 5 min with 2 mM ATPγS (lithium salt, Sigma Aldrich). 4 µL of this mixture were then applied to the plasma-cleaned grids, which were blotted for 4-6 sec using a Vitrobot Mk IV (Thermo Fisher Scientific) and vitrified by plunging into liquid ethane at liquid N_2_ temperature. Specimens for CAK-MAT1Δ219-THZ1 were prepared analogously, except that β-mercaptoethanol and ATPγS were omitted from the dilution buffer, and that CAK was diluted only 5x from the frozen stock solution. Grids were initially screened using a Tecnai F20 cryo-electron microscope (Thermo Fisher Scientific/FEI) equipped with a Gatan US 4000 CCD camera, and promising specimens after optimization of conditions were clipped into autoloader cartridges.

### Cryo-EM data collection

All datasets were acquired using a Thermo Fisher Scientific Talos Arctica cryo-EM operated at 200 kV acceleration voltage according to conventional cryo-EM approach (Herzik et al., 2019). Electron micrograph movies (example shown in Fig. S1D) were recorded using a K3 camera (Gatan) operated in super-resolution counting mode, with the microscope set to 69,444 x magnification (0.72 Å/physical pixel on the camera) at 35 e^-^/Å^2^ per second for a total dose of 69 e^-^/Å^2^ during a 2-second movie exposure, which was fractionated into 69 frames. Defocus values ranged from approx. 0.3 μm to 2.5 μm. Data collection was controlled by SerialEM (Schorb et al., 2019), using image shift for rapid image acquisition in a 3 × 3 pattern in 9 holes with active beam tilt compensation enabled.

The structure of the CAK complex was obtained from two data collection sessions. One session used a specimen in which a Pol II-CTD peptide mimic had been fused to the N-terminus of CDK7; however, no density for the peptide was observed and the structures of the two complexes were identical. The data were thus merged with the wild-type CAK data to obtain a higher-resolution reconstruction (Fig. S2A).

### Cryo-EM data processing

Cryo-EM data were processed in RELION 3.1 (Zivanov et al., 2018) as detailed in Fig. S2 and S8. Movie frames were aligned using MOTIONCOR2 (Zheng et al., 2017) from within RELION 3.1 and binned 2x (to 0.72 Å/pixel) during motion correction. After defocus estimation and contrast transfer function (CTF) fitting using CTFFIND4 (Rohou & Grigorieff, 2015) from within RELION, particles were selected using the 3D-reference-based autopicking routine in RELION 3.1. To obtain the initial autopicking template, particles were selected using the Laplacian-of-Gaussian (LoG) algorithm in RELION 3.1 (Zivanov et al., 2018) from a preliminary dataset, which was not included in this manuscript because the dataset quality compared unfavorably to the subsequent datasets used here. To obtain an initial cryo-EM reconstruction, these particles were 2D-classified and aligned to an initial reference model generated from converting PDB coordinates of CDK7 and cyclin H (PDB ID 1UA2, 1KXU) to a map using UCSF Chimera (Goddard et al., 2007). The cryo-EM map refined from these LoG-picked particles subsequently served as the autopicking template and initial reference for processing of all other datasets.

From a typical micrograph with high CAK concentration, 1,000-1,500 particles could be picked. To speed up calculations given the computational cost of processing several million particle picks per dataset, particles were initially extracted at 2.88 Å/pixel in 64-pixel boxes for initial 2D and 3D classification. Initial 2D classifications were performed using fast subsets in RELION 3.1 (Zivanov et al., 2018) and the resulting classes were generously selected, excluding mostly false positives or ice contamination. The subsequent low-resolution refinement reached near-Nyquist frequency resolutions, and 3D classification without alignment (which applies to all 3D classifications in this work) from this refinement allowed selection of a pool of high-quality particles that immediately refined to better than 4-Å resolution upon re-extractions of the particles at 1.28 Å/pixel (Fig. S2A). These particles were then either directly subjected to Bayesian polishing (Zivanov et al., 2019) in RELION 3.1 (dataset 2) or subclassified and polished (dataset 1). In subclassifications, class selection was guided mostly by inspection of the slices of the 3D volumes and classes with the crispest features were retained. During polishing, the box size was expanded from 144 pixels to 216 pixels to account for signal delocalization by the CTF (Sigworth, 2016). In our case, the resolution gain from this procedure was minimal, possibly because the high-resolution signal is contributed mostly by very low defocus particles, which are less affected by signal delocalization. However, more particles were retained in equivalent 3D classification steps than when using smaller boxes, which appeared to benefit the map quality, and thus the larger box sizes were retained. After a 2D classification to remove particles impacted by detector defects and other artifacts revealed by polishing and box size expansion, these datasets refined to resolution of 3.2-3.3 Å. One more round of 3D classification and optimized CTF refinement (Zivanov et al., 2018; Zivanov et al., 2020) yielded subsets of 67,890 and 101,016 particles, which were joined, subjected to CTF refinement again, and refined to the final 2.9-Å resolution map after removal of all particles that came from micrographs with power spectra that did not extend to 4 Å or better.

The resolution of the reconstruction, obtained by gold-standard refinement of fully independent particle subsets (Scheres & Chen, 2012), was assessed by Fourier Shell Correlation (FSC) (Fig. S2B) at the FSC = 0.143 cut-off (Rosenthal & Henderson, 2003). Additionally, the 3D reconstruction was validated using the 3D FSC validation server (Tan et al., 2017), which confirmed that the map is isotropic (Fig. S2C) with a sphericity of 0.96.

The procedure for reconstruction of the CAK-THZ1 structure (Fig. S8A) was essentially as described for the ATPγS-bound reconstruction, with minor modifications. Apparent larger counts of low-quality particle images necessitated two 3D classification steps at the 2.88 Å/pixel stage to eliminate these and reach 5.9 Å resolution before particle re-extraction at smaller pixel size. Due to the lower overall resolution, the final pixel size was chosen to be 1.44 Å/pixel, and the corresponding enlarged box size after particle polishing was 192 pixels. The resolution of the map immediately after polishing (from 310,052 particles) was estimated at 3.3 Å, which is nominally better than the 3.4 Å estimated for the final map from 31,198 particles, obtained after two more 3D classification steps. However, the latter map exhibited far better side chain density, better overall connectivity, and a lower B-factor (approx. -60 Å^2^ vs. -90 Å^2^ as automatically determined by the RELION post processing function) and was therefore used for interpretation, model building, and refinement.

### Model building, refinement, and validation

Before model building, the 3D volumes obtained from cryo-EM map refinement were re-boxed to 144 pixels (CAK reconstruction) or 128 pixels (CAK-THZ1) without changing the pixel size. The atomic model of the CAK complex was assembled by fitting the high-resolution structures of CDK7 and cyclin H (Andersen et al., 1997; Kim et al., 1996; Lolli et al., 2004) into the cryo-EM map using UCSF Chimera (Goddard et al., 2007), followed by re-building in COOT (Emsley et al., 2010). Both crystal structures showed a good initial fit to the cryo-EM map and needed only minor rebuilding, with the exception of the CDK7 T-loop, which occupies the inactive conformation in the crystal structure, and CDK7 residues 104-111, which were found to be register-shifted by one residue. After rebuilding of CDK7 and cyclin H according to the density, a continuous stretch of remaining unassigned density was identified and assigned to MAT1 based on agreement of the density with secondary structure prediction and the pattern of large side chains near the C-terminus of MAT1 (Fig. 1C). The model was then iteratively rebuilt in COOT and refined using the real space refinement program in PHENIX (Adams et al., 2010; Afonine et al., 2018). To maintain good model geometry at the given resolution, Ramachandran, rotamer, Cβ, and secondary structure restraints were used throughout.

The molecular structure and geometric restraints for THZ1 were generated using PHENIX ELBOW (Moriarty et al., 2009), and the CAK-THZ1 structure was initially refined using the higher-resolution CAK-ATPγS structure to provide reference restraints, before 5 final macro-cycles of real-space refinement were carried out without reference restraints.

The structures were validated by computing the FSC between the model and the map to assess the agreement of the model and the cryo-EM density (Fig. S2B, S8B), and using MOLPROBITY (Williams et al., 2018) to assess the quality of the model geometry. Validation statistics are given in Table S1. Components of the refined atomic models are listed in Table S2.

### Mass spectrometry

10-µL aliquots of CAK and CAK-THZ1 were diluted to an approximate concentration of 10 µM and subjected to mass spectrometry analysis to evaluate the binding of THZ1 to the CAK. Intact protein samples were analyzed using a Synapt G2-S*i* mass spectrometer (Waters, Milford, MA), as described (Light et al., 2018). The deconvoluted mass spectra for the samples are shown in Fig. S7.

### Other software used

Software used in this project was configured by SBGRID (Morin et al., 2013). Molecular graphics were created using UCSF Chimera (Goddard et al., 2007), UCSF ChimeraX (Goddard et al., 2018), and PyMol (The PyMol molecular graphics system, Schrödinger LLC). Local resolution of cryo-EM maps was computed using RELION 3.1 (Zivanov et al., 2018). Buried surface areas were computed using PDBePISA (Krissinel & Henrick, 2007).

## Acknowledgments

We thank A. Chintangal and P. Tobias for support with computation, J. Remis and P. Grob for support with microscopy, K. Dong and A. Martin for help with Western blot imaging, and A. Killilea and M. Fischer at the UC Berkeley cell culture facility for providing insect cell cultures. This work was funded through NIGMS grants R01-GM63072, P01-GM063210, and R35-GM127018 to E.N. The Synapt mass spectrometer was purchased with NIH support (grant 1S10OD020062-01). E.N. is a Howard Hughes Medical Institute Investigator.

## Accession Codes

Cryo-EM maps of CAK and CAK-MAT1Δ219-THZ1 have been deposited at the EM Data Resource with accession codes EMD-XXXX and EMD-XXXX. The refined coordinate models have been deposited at the Protein Data Bank (PDB) with accession codes XXXX and XXXX, respectively.

## References

Adams P. D., Afonine P. V., Bunkóczi G., Chen V. B., Davis I. W., Echols N., Headd J. J., Hung L. W., Kapral G. J., Grosse-Kunstleve R. W., McCoy A. J., Moriarty N. W., Oeffner R., Read R. J., Richardson D. C., Richardson J. S., Terwilliger T. C., Zwart P. H. (2010) PHENIX: a comprehensive Python-based system for macromolecular structure solution. Acta Crystallogr D Biol Crystallogr 66 (Pt 2): 213–221

Afonine P. V., Poon B. K., Read R. J., Sobolev O. V., Terwilliger T. C., Urzhumtsev A., Adams P. D. (2018) Real-space refinement in PHENIX for cryo-EM and crystallography. Acta Crystallogr D Struct Biol 74 (Pt 6): 531–544

Akoulitchev S., Reinberg D. (1998) The molecular mechanism of mitotic inhibition of TFIIH is mediated by phosphorylation of CDK7. Genes & Dev 12(22): 3541–3550

Andersen G., Busso D., Poterszman A., Hwang J. R., Wurtz J. M., Ripp R., Thierry J. C., Egly J. M., Moras D. (1997) The structure of cyclin H: common mode of kinase activation and specific features. EMBO J 16(5): 958–967

Baumli S., Lolli G., Lowe E. D., Troiani S., Rusconi L., Bullock A. N., Debreczeni J. E., Knapp S., Johnson L. N. (2008) The structure of P-TEFb (CDK9/cyclin T1), its complex with flavopiridol and regulation by phosphorylation. EMBO J 27(13): 1907–1918

Bieniossek C., Richmond T. J., Berger I. (2008) MultiBac: multigene baculovirus-based eukaryotic protein complex production. Curr Protoc Protein Sci Chapter 5 Unit 5.20

Brown N. R., Lowe E. D., Petri E., Skamnaki V., Antrobus R., Johnson L. N. (2007) Cyclin B and cyclin A confer different substrate recognition properties on CDK2. Cell Cycle 6(11): 1350–1359

Busso D., Keriel A., Sandrock B., Poterszman A., Gileadi O., Egly J. M. (2000) Distinct regions of MAT1 regulate cdk7 kinase and TFIIH transcription activities. J Biol Chem 275(30): 22815–22823

Connell-Crowley L., Solomon M. J., Wei N., Harper J. W. (1993) Phosphorylation independent activation of human cyclin-dependent kinase 2 by cyclin A in vitro. Mol Biol Cell 4(1): 79–92

Devault A., Martinez A. M., Fesquet D., Labbé J. C., Morin N., Tassan J. P., Nigg E. A., Cavadore J. C., Dorée M. (1995) MAT1 (‘menage à trois’) a new RING finger protein subunit stabilizing cyclin H-cdk7 complexes in starfish and Xenopus CAK. EMBO J 14(20): 5027–5036

Echalier A., Endicott J. A., Noble M. E. M. (2010) Recent developments in cyclin-dependent kinase biochemical and structural studies. Biochimica et biophysica acta 1804(3): 511–519

Emsley P., Lohkamp B., Scott W. G., Cowtan K. (2010) Features and development of Coot. Acta Crystallogr D Biol Crystallogr 66 (Pt 4): 486–501

Feaver W. J., Svejstrup J. Q., Henry N. L., Kornberg R. D. (1994) Relationship of CDK-activating kinase and RNA polymerase II CTD kinase TFIIH/TFIIK. Cell 79(6): 1103–1109

Fisher R. P., Jin P., Chamberlin H. M., Morgan D. O. (1995) Alternative mechanisms of CAK assembly require an assembly factor or an activating kinase. Cell 83(1): 47–57

Fisher R. P., Morgan D. O. (1994) A novel cyclin associates with MO15/CDK7 to form the CDK-activating kinase. Cell 78(4): 713–724

Galbraith M. D., Bender H., Espinosa J. M. (2019) Therapeutic targeting of transcriptional cyclin-dependent kinases. Transcription 10(2): 118–136

Goddard T. D., Huang C. C., Ferrin T. E. (2007) Visualizing density maps with UCSF Chimera. J Struct Biol 157(1): 281–287

Goddard T. D., Huang C. C., Meng E. C., Pettersen E. F., Couch G. S., Morris J. H., Ferrin T. E. (2018) UCSF ChimeraX: Meeting modern challenges in visualization and analysis. Protein Sci 27(1): 14–25

Gradia S. D., Ishida J. P., Tsai M.-S., Jeans C., Tainer J. A., Fuss J. O. (2017) MacroBac: New Technologies for Robust and Efficient Large-Scale Production of Recombinant Multiprotein Complexes. Meth Enzymol 592 1–26

Greber B. J., Nguyen T. H. D., Fang J., Afonine P. V., Adams P. D., Nogales E. (2017) The cryo-electron microscopy structure of human transcription factor IIH. Nature 549(7672): 414–417

Greber B. J., Toso D., Fang J., Nogales E. (2019) The complete structure of the human TFIIH core complex. eLife 8 e44771

Harlen K. M., Churchman L. S. (2017) The code and beyond: transcription regulation by the RNA polymerase II carboxy-terminal domain. Nat Rev Mol Cell Biol 18(4): 263–273

He Y., Yan C., Fang J., Inouye C., Tjian R., Ivanov I., Nogales E. (2016) Near-atomic resolution visualization of human transcription promoter opening. Nature 533(7603): 359–365

Herzik M. A., Wu M., Lander G. C. (2019) High-resolution structure determination of sub-100 kDa complexes using conventional cryo-EM. Nat Commun 10(1): 1032

Honda R., Lowe E. D., Dubinina E., Skamnaki V., Cook A., Brown N. R., Johnson L. N. (2005) The structure of cyclin E1/CDK2: implications for CDK2 activation and CDK2-independent roles. EMBO J 24(3): 452–463

Hu S., Marineau J. J., Rajagopal N., Hamman K. B., Choi Y. J., Schmidt D. R., Ke N., Johannessen L., Bradley M. J., Orlando D. A., Alnemy S. R., Ren Y., Ciblat S., Winter D. K., Kabro A., Sprott K. T., Hodgson J. G., Fritz C. C., Carulli J. P., di Tomaso E., Olson E. R. (2019) Discovery and Characterization of SY-1365, a Selective, Covalent Inhibitor of CDK7. Cancer Res 79(13): 3479–3491

Irwin C. R., Farmer A., Willer D. O., Evans D. H. (2012) In-fusion® cloning with vaccinia virus DNA polymerase. Methods Mol Biol 890 (Chapter 2): 23–35

Jeffrey P. D., Russo A. A., Polyak K., Gibbs E., Hurwitz J., Massagué J., Pavletich N. P. (1995) Mechanism of CDK activation revealed by the structure of a cyclinA-CDK2 complex. Nature 376(6538): 313–320

Jeronimo C., Bataille A. R., Robert F. (2013) The writers, readers, and functions of the RNA polymerase II C-terminal domain code. Chem Rev 113(11): 8491–8522

Keogh M.-C., Cho E.-J., Podolny V., Buratowski S. (2002) Kin28 is found within TFIIH and a Kin28-Ccl1-Tfb3 trimer complex with differential sensitivities to T-loop phosphorylation. Mol Cell Biol 22(5): 1288–1297

Kim K. K., Chamberlin H. M., Morgan D. O., Kim S. H. (1996) Three-dimensional structure of human cyclin H, a positive regulator of the CDK-activating kinase. Nat Struct Biol 3(10): 849–855

Kokic G., Chernev A., Tegunov D., Dienemann C., Urlaub H., Cramer P. (2019) Structural basis of TFIIH activation for nucleotide excision repair. Nat Commun 10(1): 2885

Krissinel E., Henrick K. (2007) Inference of macromolecular assemblies from crystalline state. J Mol Biol 372(3): 774–797

Kwiatkowski N., Zhang T., Rahl P. B., Abraham B. J., Reddy J., Ficarro S. B., Dastur A., Amzallag A., Ramaswamy S., Tesar B., Jenkins C. E., Hannett N. M., McMillin D., Sanda T., Sim T., Kim N. D., Look T., Mitsiades C. S., Weng A. P., Brown J. R., Benes C. H., Marto J. A., Young R. A., Gray N. S. (2014) Targeting transcription regulation in cancer with a covalent CDK7 inhibitor. Nature 511(7511): 616–620

Labbé J. C., Martinez A. M., Fesquet D., Capony J. P., Darbon J. M., Derancourt J., Devault A., Morin N., Cavadore J. C., Dorée M. (1994) p40MO15 associates with a p36 subunit and requires both nuclear translocation and Thr176 phosphorylation to generate cdk-activating kinase activity in Xenopus oocytes. EMBO J 13(21): 5155–5164

Larochelle S., Chen J., Knights R., Pandur J., Morcillo P., Erdjument-Bromage H., Tempst P., Suter B., Fisher R. P. (2001) T-loop phosphorylation stabilizes the CDK7-cyclin H-MAT1 complex in vivo and regulates its CTD kinase activity. EMBO J 20(14): 3749–3759

Larochelle S., Pandur J., Fisher R. P., Salz H. K., Suter B. (1998) Cdk7 is essential for mitosis and for in vivo Cdk-activating kinase activity. Genes Dev 12(3): 370–381

Li M. Z., Elledge S. J. (2012) SLIC: a method for sequence- and ligation-independent cloning. Methods Mol Biol 852 51–59

Light S. H., Su L., Rivera-Lugo R., Cornejo J. A., Louie A., Iavarone A. T., Ajo-Franklin C. M., Portnoy D. A. (2018) A flavin-based extracellular electron transfer mechanism in diverse Gram-positive bacteria. Nature 562(7725): 140–144

Lolli G., Lowe E. D., Brown N. R., Johnson L. N. (2004) The crystal structure of human CDK7 and its protein recognition properties. Structure 12(11): 2067–2079

Ma Z., Wu Y., Jin J., Yan J., Kuang S., Zhou M., Zhang Y., Guo A.-Y. (2013) Phylogenetic analysis reveals the evolution and diversification of cyclins in eukaryotes. Mol Phylogenet Evol 66(3): 1002–1010

Martinez A. M., Afshar M., Martin F., Cavadore J. C., Labbé J. C., Dorée M. (1997) Dual phosphorylation of the T-loop in cdk7: its role in controlling cyclin H binding and CAK activity. EMBO J 16(2): 343–354

Morgan D. O. (1997) Cyclin-dependent kinases: engines, clocks, and microprocessors. Annu Rev Cell Dev Biol 13 261–291

Moriarty N. W., Grosse-Kunstleve R. W., Adams P. D. (2009) electronic Ligand Builder and Optimization Workbench (eLBOW): a tool for ligand coordinate and restraint generation. Acta Crystallogr D Biol Crystallogr 65 (Pt 10): 1074–1080

Morin A., Eisenbraun B., Key J., Sanschagrin P. C., Timony M. A., Ottaviano M., Sliz P. (2013) Collaboration gets the most out of software. eLife 2 e01456

Myers L. C., Gustafsson C. M., Bushnell D. A., Lui M., Erdjument-Bromage H., Tempst P., Kornberg R. D. (1998) The Med proteins of yeast and their function through the RNA polymerase II carboxy-terminal domain. Genes Dev 12(1): 45–54

Peissert S., Sauer F., Grabarczyk D. B., Braun C., Sander G., Poterszman A., Egly J.-M., Kuper J., Kisker C. (2020) In TFIIH the Arch domain of XPD is mechanistically essential for transcription and DNA repair. Nat Commun 11(1): 1667–1613

Rohou A., Grigorieff N. (2015) CTFFIND4: Fast and accurate defocus estimation from electron micrographs. J Struct Biol 192(2): 216–221

Rosenthal P. B., Henderson R. (2003) Optimal determination of particle orientation, absolute hand, and contrast loss in single-particle electron cryomicroscopy. J Mol Biol 333(4): 721–745

Russo A. A., Jeffrey P. D., Pavletich N. P. (1996) Structural basis of cyclin-dependent kinase activation by phosphorylation. Nat Struct Biol 3(8): 696–700

Sainsbury S., Bernecky C., Cramer P. (2015) Structural basis of transcription initiation by RNA polymerase II. Nat Rev Mol Cell Biol 16(3): 129–143

Scheres S. H. W., Chen S. (2012) Prevention of overfitting in cryo-EM structure determination. Nat Meth 9(9): 853–854

Schilbach S., Hantsche M., Tegunov D., Dienemann C., Wigge C., Urlaub H., Cramer P. (2017) Structures of transcription pre-initiation complex with TFIIH and Mediator. Nature 551(7679): 204–209

Schorb M., Haberbosch I., Hagen W. J. H., Schwab Y., Mastronarde D. N. (2019) Software tools for automated transmission electron microscopy. Nat Meth 16(6): 471–477

Serizawa H., Mäkelä T. P., Conaway J. W., Conaway R. C., Weinberg R. A., Young R. A. (1995) Association of Cdk-activating kinase subunits with transcription factor TFIIH. Nature 374(6519): 280–282

Shiekhattar R., Mermelstein F., Fisher R. P., Drapkin R., Dynlacht B., Wessling H. C., Morgan D. O., Reinberg D. (1995) Cdk-activating kinase complex is a component of human transcription factor TFIIH. Nature 374(6519): 283–287

Sigworth F. J. (2016) Principles of cryo-EM single-particle image processing. Microscopy (Tokyo) 65(1): 57–67

Søgaard T. M. M., Svejstrup J. Q. (2007) Hyperphosphorylation of the C-terminal repeat domain of RNA polymerase II facilitates dissociation of its complex with mediator. J Biol Chem 282(19): 14113–14120

Tan Y. Z., Baldwin P. R., Davis J. H., Williamson J. R., Potter C. S., Carragher B., Lyumkis D. (2017) Addressing preferred specimen orientation in single-particle cryo-EM through tilting. Nat Meth 14(8): 793–796

Tassan J. P., Jaquenoud M., Fry A. M., Frutiger S., Hughes G. J., Nigg E. A. (1995) In vitro assembly of a functional human CDK7-cyclin H complex requires MAT1, a novel 36 kDa RING finger protein. EMBO J 14(22): 5608–5617

Williams C. J., Headd J. J., Moriarty N. W., Prisant M. G., Videau L. L., Deis L. N., Verma V., Keedy D. A., Hintze B. J., Chen V. B., Jain S., Lewis S. M., Arendall W. B., Snoeyink J., Adams P. D., Lovell S. C., Richardson J. S., Richardson D. C. (2018) MolProbity: More and better reference data for improved all-atom structure validation. Protein Sci 27(1): 293–315

Wood D. J., Endicott J. A. (2018) Structural insights into the functional diversity of the CDK-cyclin family. Open biology 8(9): 180112–180126

Yan C., Dodd T., He Y., Tainer J. A., Tsutakawa S. E., Ivanov I. (2019) Transcription preinitiation complex structure and dynamics provide insight into genetic diseases. Nat Struct Mol Biol 26(6): 397–406

Yee A., Nichols M. A., Wu L., Hall F. L., Kobayashi R., Xiong Y. (1995) Molecular cloning of CDK7-associated human MAT1, a cyclin-dependent kinase-activating kinase (CAK) assembly factor. Cancer Res 55(24): 6058–6062

Zheng S. Q., Palovcak E., Armache J.-P., Verba K. A., Cheng Y., Agard D. A. (2017) MotionCor2: anisotropic correction of beam-induced motion for improved cryo-electron microscopy. Nat Meth 14(4): 331–332

Zivanov J., Nakane T., Forsberg B. O., Kimanius D., Hagen W. J., Lindahl E., Scheres S. H. (2018) New tools for automated high-resolution cryo-EM structure determination in RELION-3. eLife 7 e42166

Zivanov J., Nakane T., Scheres S. H. W. (2019) A Bayesian approach to beam-induced motion correction in cryo-EM single-particle analysis. IUCrJ 6 (Pt 1): 5-17

Zivanov J., Nakane T., Scheres S. H. W. (2020) Estimation of high-order aberrations and anisotropic magnification from cryo-EM data sets in RELION-3.1. IUCrJ 7 (Pt 2): 253–267

