## Supplementary Information PDF for "The cryo-electron microscopy structure of the human CDK-activating kinase"

**This PDF file includes:**

Figures S1 to S8
Tables S1 and S2
SI References

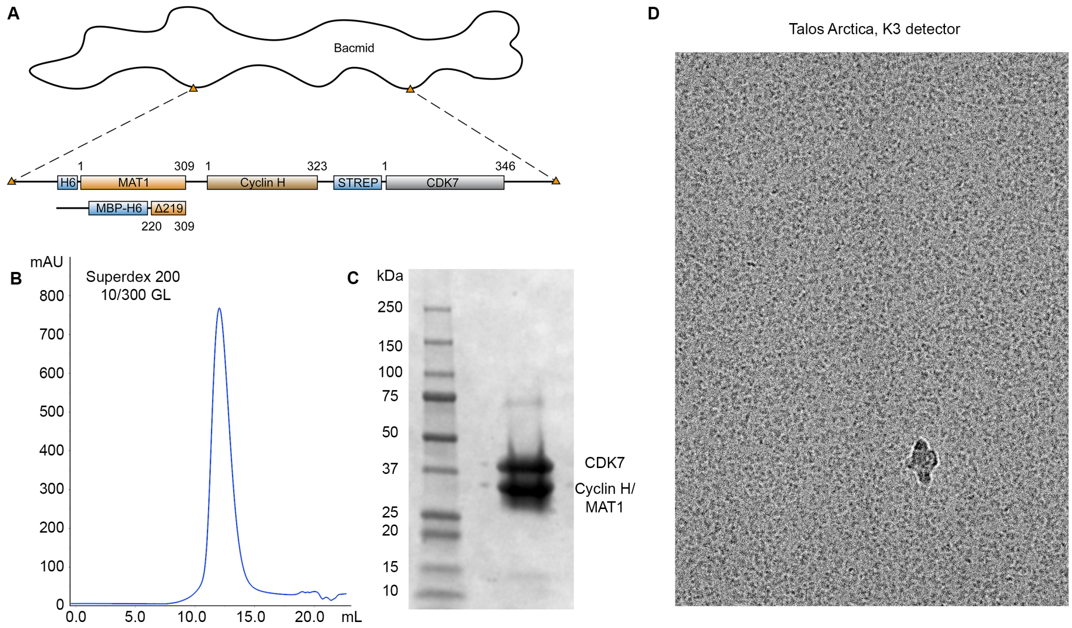


**Figure S1. CAK expression, purification, and specimen preparation**. (**A**) Schematic of baculovirus-based expression construct used for expression of full-length CAK and CAK-MAT1Δ219 in insect cells. (**B**) Elution profile from Superdex 200 10/300 GL column showing a monodisperse peak for full-length CAK. (**C**) SDS-PAGE gel of purified CAK. The bands for cyclin H and MAT1 overlap due to their similar molecular weights. (**D**) Representative electron micrograph showing CAK particles, acquired on a 200 kV-Talos Arctica electron microscope using a K3 direct electron detector.

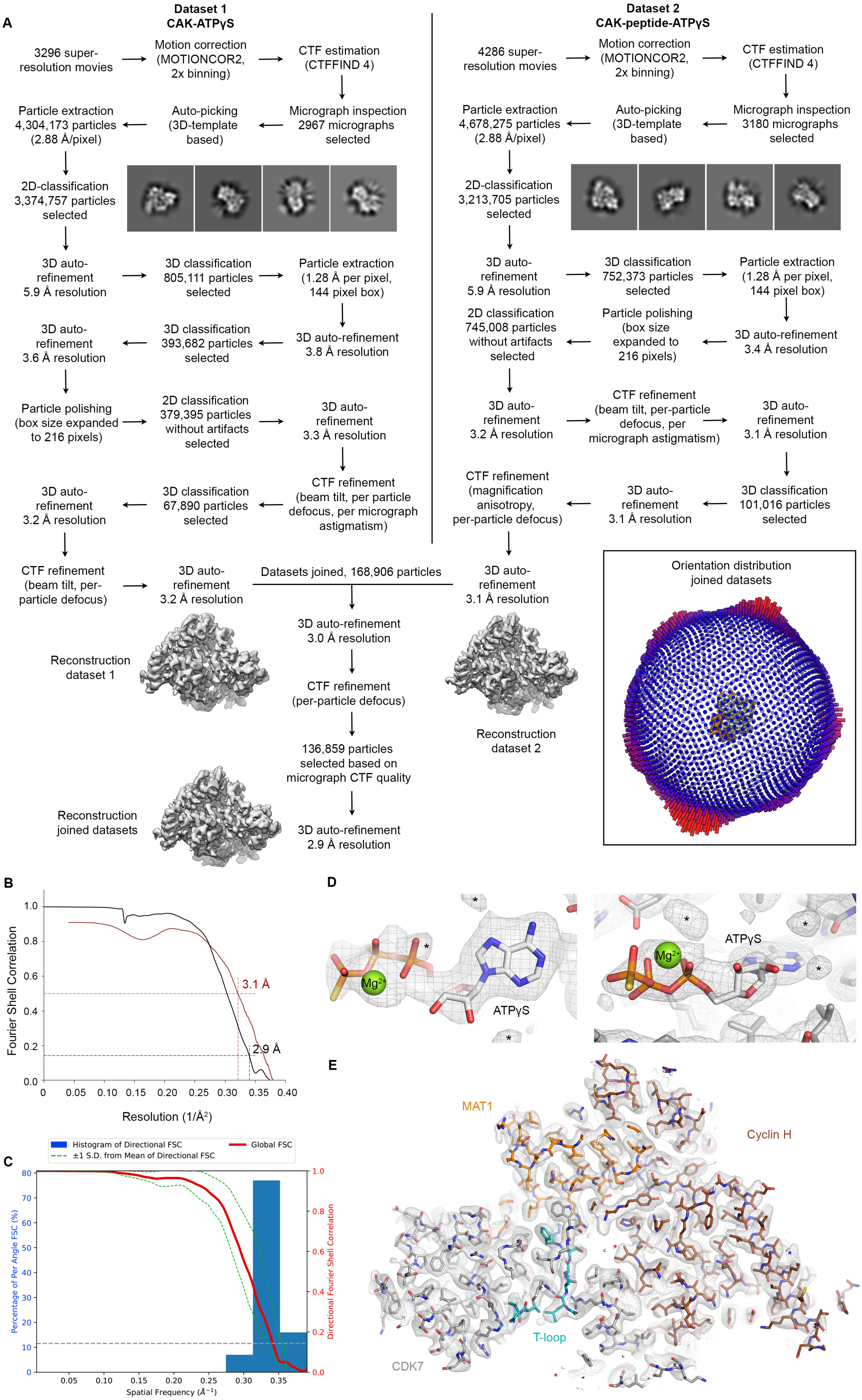

A
Dataset 1
CAK-ATPγS
3296 super-resolution movies
Motion correction (MOTIONCOR2, 2x binning)
CTF estimation (CTFFIND 4)
Micrograph inspection 2967 micrographs selected
Auto-picking (3D-template based)
Particle extraction 4,304,173 particles (2.88 Å/pixel)
2D-classification 3,374,757 particles selected
3D auto-refinement 5.9 Å resolution
3D classification 805,111 particles selected
Particle extraction (1.28 Å per pixel, 144 pixel box)
3D auto-refinement 3.8 Å resolution
3D classification 393,682 particles selected
3D auto-refinement 3.6 Å resolution
Particle polishing (box size expanded to 216 pixels)
2D classification 379,395 particles without artifacts selected
3D auto-refinement 3.3 Å resolution
CTF refinement (beam tilt, per particle defocus, per micrograph astigmatism)
3D classification 67,890 particles selected
3D auto-refinement 3.2 Å resolution
CTF refinement (beam tilt, per-particle defocus)
3D auto-refinement 3.2 Å resolution
Reconstruction dataset 1
Dataset 2
CAK-peptide-ATPγS
4286 super-resolution movies
Motion correction (MOTIONCOR2, 2x binning)
CTF estimation (CTFFIND 4)
Micrograph inspection 3180 micrographs selected
Auto-picking (3D-template based)
Particle extraction 4,678,275 particles (2.88 Å/pixel)
2D-classification 3,213,705 particles selected
3D auto-refinement 5.9 Å resolution
3D classification 752,373 particles selected
Particle extraction (1.28 Å per pixel, 144 pixel box)
3D auto-refinement 3.4 Å resolution
Particle polishing (box size expanded to 216 pixels)
2D classification 745,008 particles without artifacts selected
3D auto-refinement 3.2 Å resolution
CTF refinement (beam tilt, per-particle defocus, per micrograph astigmatism)
3D auto-refinement 3.1 Å resolution
3D classification 101,016 particles selected
3D auto-refinement 3.1 Å resolution
CTF refinement (magnification anisotropy, per-particle defocus)
3D auto-refinement 3.1 Å resolution
Reconstruction dataset 2
Datasets joined, 168,906 particles
3D auto-refinement 3.0 Å resolution
CTF refinement (per-particle defocus)
136,859 particles selected based on micrograph CTF quality
3D auto-refinement 2.9 Å resolution
Reconstruction joined datasets
Orientation distribution joined datasets
B
Fourier Shell Correlation
Resolution (1/Å²)
3.1 Å
2.9 Å
C
Histogram of Directional FSC
±1 S.D. from Mean of Directional FSC
Global FSC
Percentage of Per Angle FSC (%)
Directional Fourier Shell Correlation
Spatial Frequency (Å⁻¹)
D
Mg²⁺
ATPγS
E
MAT1
Cyclin H
T-loop
CDK7

**Figure S2. Structure determination of the CAK-ATPγS complex.** (**A**) Data processing scheme. Due to the large number of operations, only selected intermediate results are shown. The strategy was analogous to the processing of the CAK-THZ1 dataset (Fig. S6), for which detailed depictions of classifications and intermediate maps are shown. Inset: Projection orientation distribution of the final refinement. CAK is preferentially oriented but shows a continuous series of views along one axis, which is sufficient to obtain complete sampling of Fourier space and reconstruction of an isotropic map (see C). (**B**) Fourier shell curves between independent cryo-EM half-maps (black) and between the refined coordinate model and the cryo-EM map (red). (**C**) Validation of the cryo-EM map of the human CAK using the 3D FSC server (Tan et al., 2017) confirms that the map contains structural information at better than approx. 3.3-Å resolution in all directions (sphericity > 0.96). (**D**) Density for ATPγS in the active site of CDK7. Additional densities near the nucleotide (indicated by asterisks) may correspond to additional metal ions or unassigned ordered solvent molecules. (**E**) Section of the cryo-EM density with the fitted model.

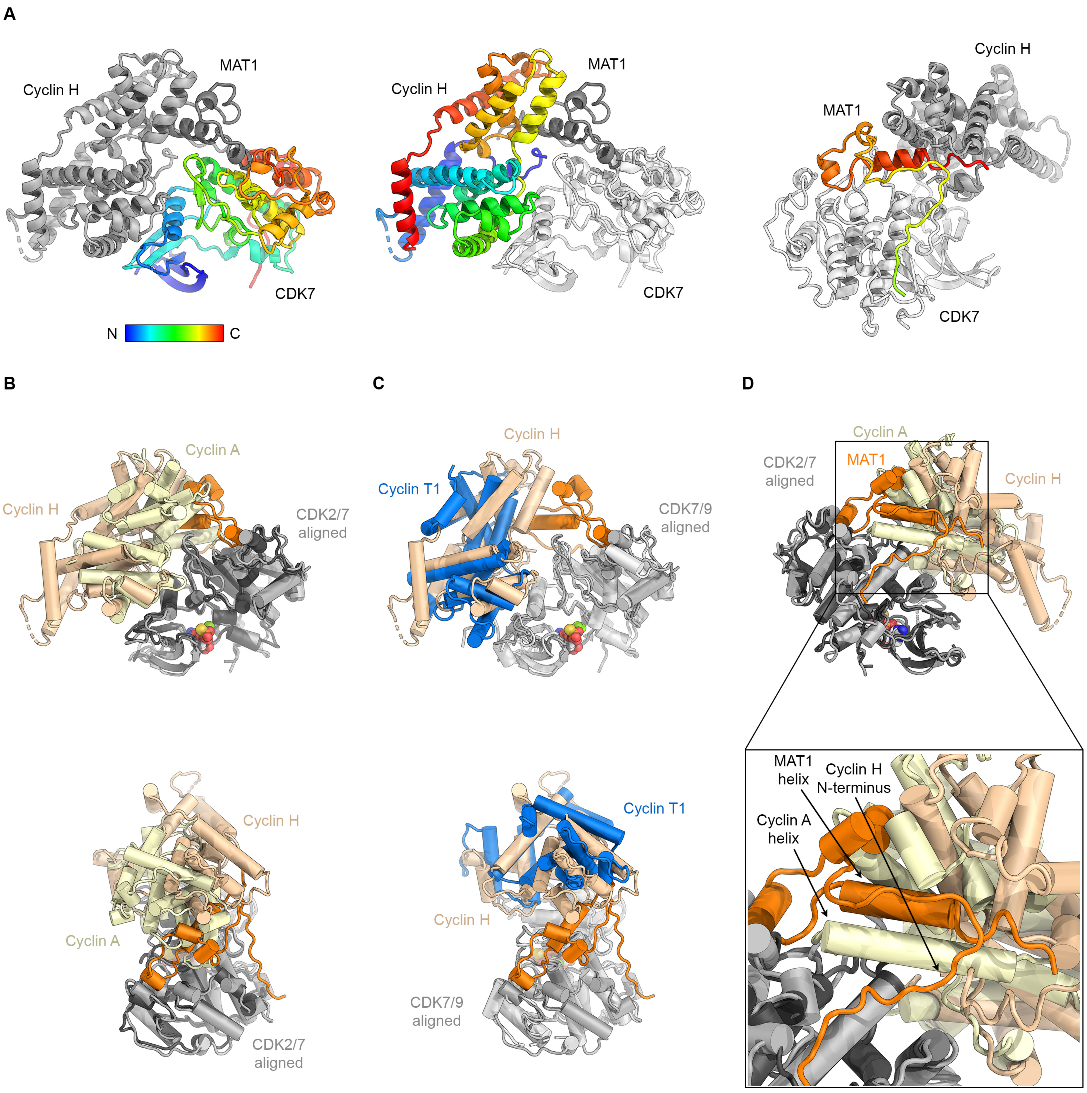


**Figure S3. Comparison to other CDK-cyclin structures.** (**A**) CDK7, cyclin H, and MAT1 colored by residue from N-terminus (blue) to C-terminus (red). (**B**) Superposition of the CAK structure with the cell-cycle controlling CDK2-cyclin A complex (PDB ID 1FIN) (Jeffrey et al., 1995). Cyclin H (light brown) in the CAK is rotated to the back slightly and shows a widened gap between the CDK and the cyclin compared to CDK2-cyclin A (yellow). MAT1 would not fit between the CDK and the cyclin in the cell-cycle CDK-like configuration. (**C**) Superposition of the CAK with the structure of the transcription-controlling complex pTEF-B (CDK9-cyclin T1; cyclin T1 in blue) (PDB ID 3BLG) (Baumli et al., 2008). Cyclin T1 is rotated away from CDK9 even further than cyclin H is from CDK7. (**D**) Cyclin A (yellow) in the CDK2-cyclin A complex (PDB ID 1FIN) (Jeffrey et al., 1995) has an α-helix that extends further from the location of the N-terminus of cyclin H (light brown). The C-terminal α-helix of MAT1 is localized similarly to this cyclin A helix and also interacts with CDK7.

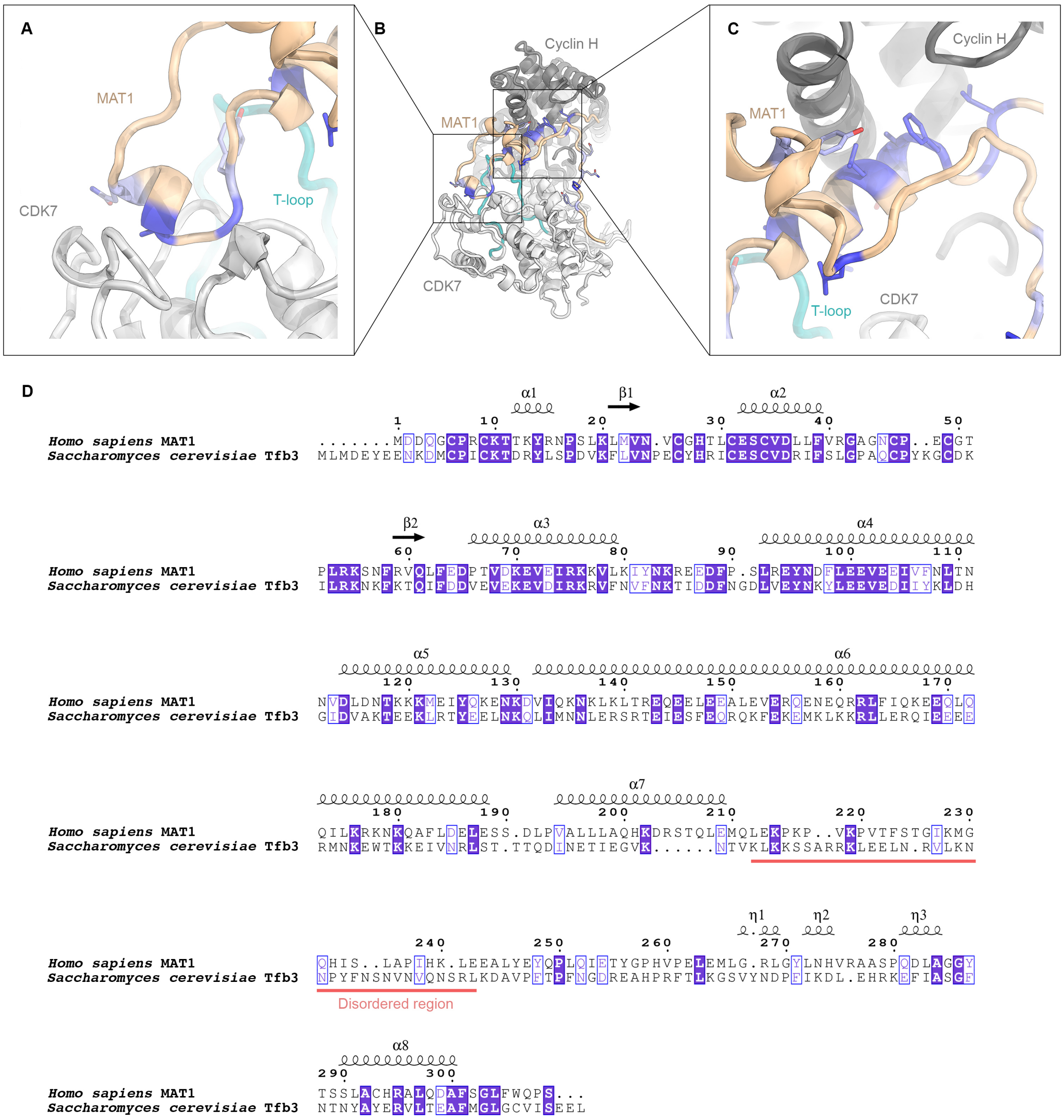


**Figure S4. Conservation of MAT1 and binding to Mediator in the Pol II-PIC.** (**A-C**) Residues that are conserved between human MAT1 and yeast Tfb3 are shown in violet (darker shade = identical; lighter shade = similar). (**D**) Sequence alignment of MAT1 and Tfb3, extracted from the multiple sequence alignment shown in Fig. S5. Residues are color-coded as in A-C.

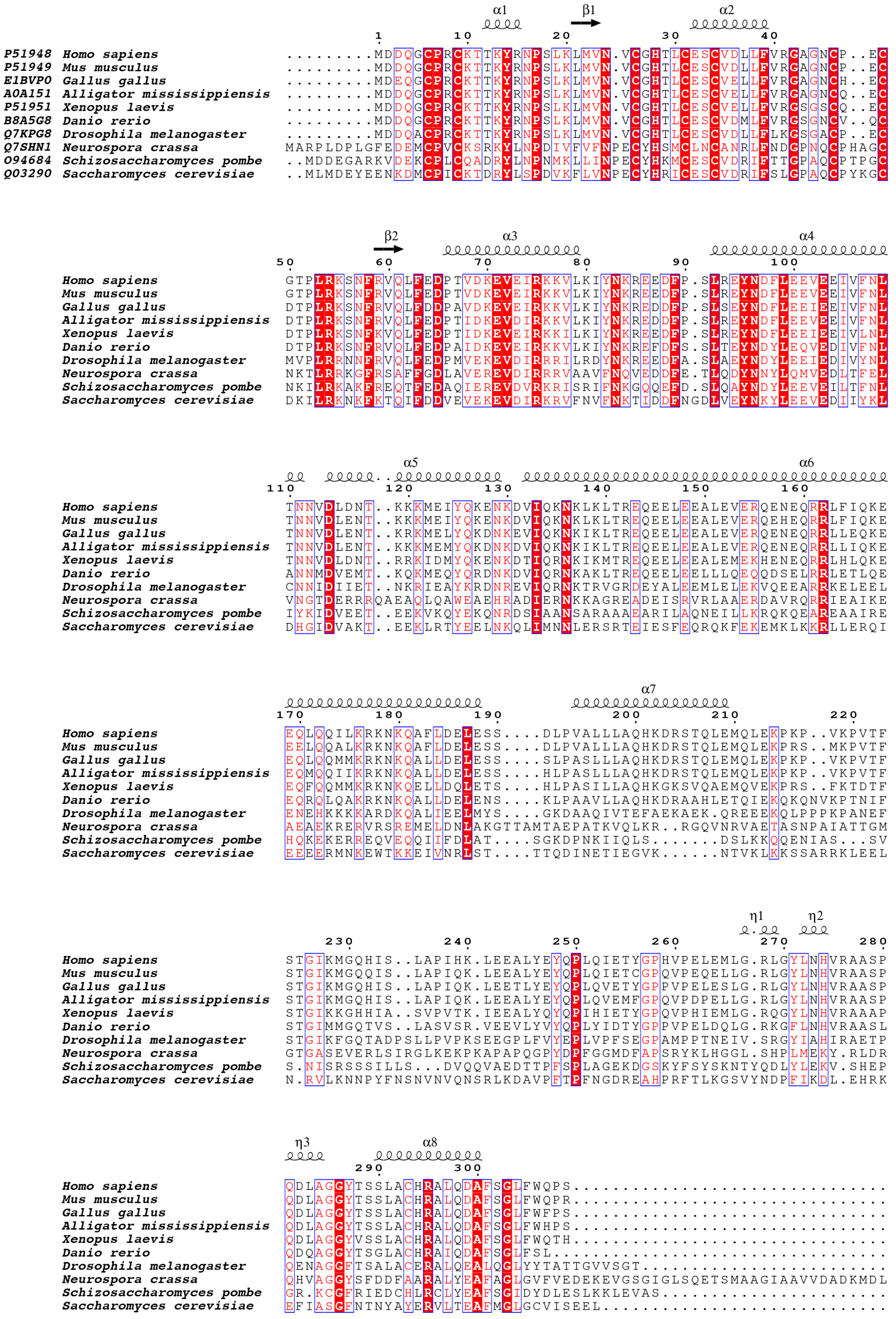


**Figure S5. Multiple sequence alignment of MAT1 from different animal and fungal species.** Species names and accession numbers are indicated. The alignment was computed with Clustal Omega (Sievers et al., 2011) and visualized using the ESPRIPT web server (Robert & Gouet, 2014).

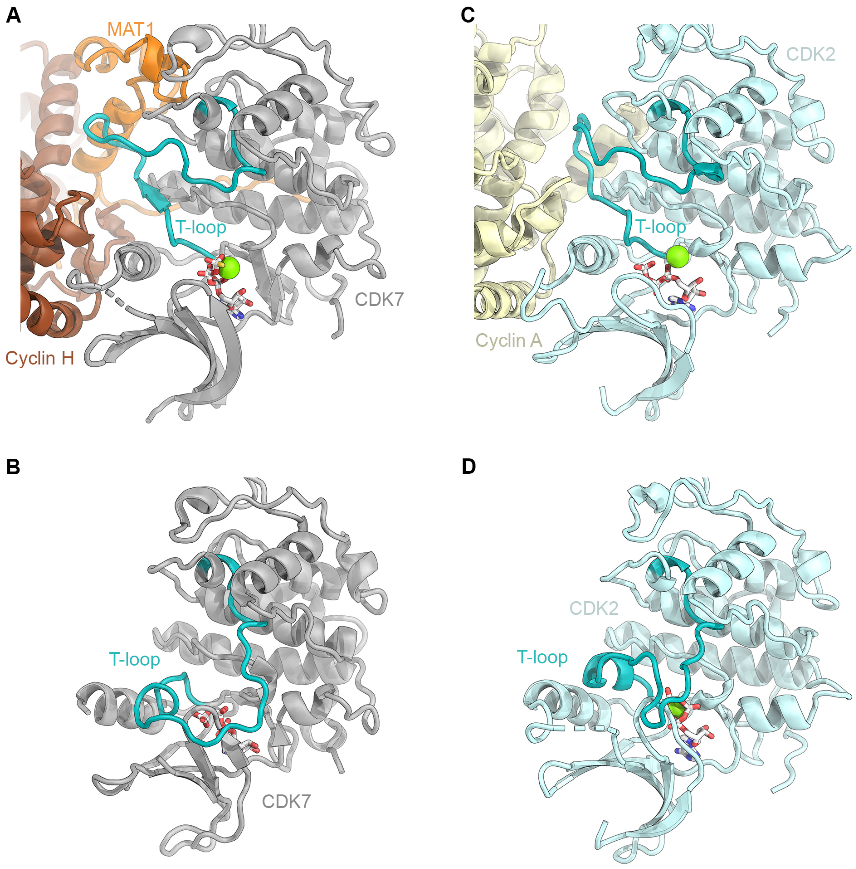


**Figure S6. CDK7 conformation and CAK activity.** (**A**) Extended conformation of the T-loop (teal) in the structure of the CAK. (**B**) The T-loop in the structure of isolated CDK7 (PDB ID 1UA2) is folded across the active site and would preclude substrate binding (Lolli et al., 2004). (**C**, **D**) T-loop conformation in the well-studied CDK2 system for comparison (Russo et al., 1996; Schulze-Gahmen et al., 1996). In the active CDK2-cyclin A complex (**C**) (PDB ID 1JST), the T-loop is extended towards cyclin A (yellow). In inactive CDK2 (**D**), the T-loop (teal) is folded across the active site (PDB ID 1HCK).

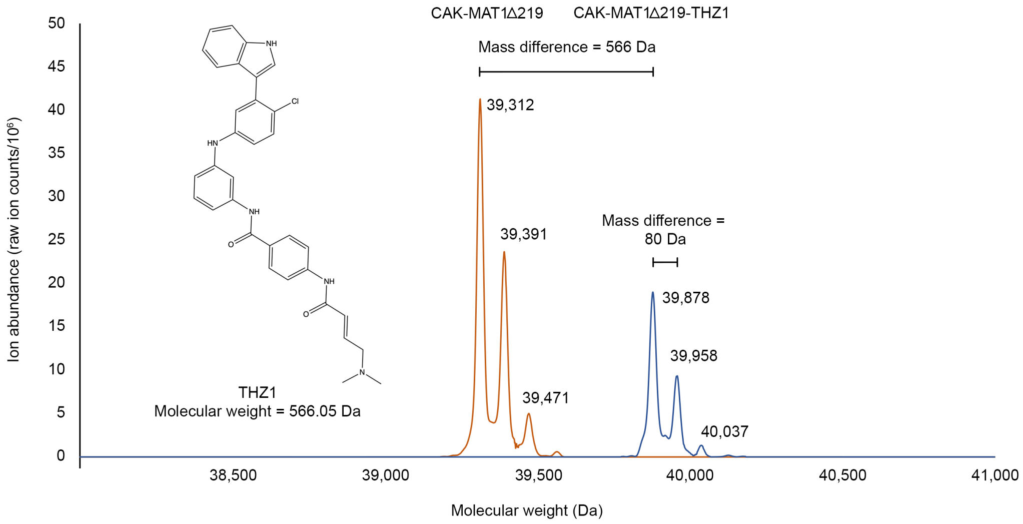


**Figure S7. Mass spectrometric analysis of THZ1 binding to CAK.** CAK-MAT1Δ219 and CAK-MAT1Δ219-THZ1 complexes were analyzed using intact electrospray ionization mass spectrometry. Spectra for CDK7 from untreated CAK are plotted in orange and spectra for CDK7 from THZ1-treated CAK are plotted in blue (cyclin H and MAT1Δ219 did not show any difference between the treated and untreated samples). A mass difference of 566 Da corresponds to the mass of THZ1 (chemical structure shown on the left). Mass differences of approx. 80 Da likely correspond to phosphorylations on CDK7.

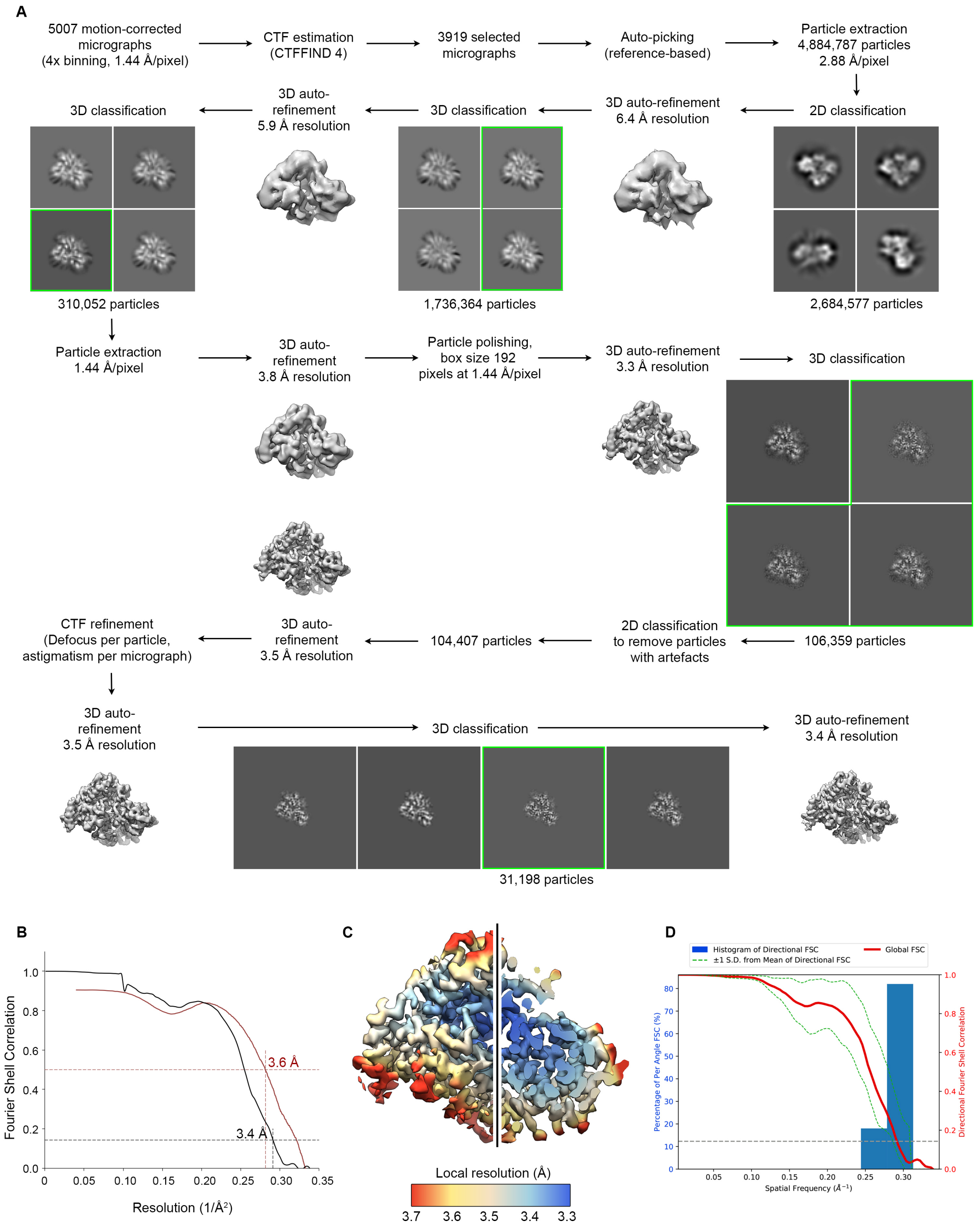


**Figure S8. Structure determination of CAK-THZ1.** (**A**) Data processing scheme. All maps shown are un-sharpened RELION outputs. (**B**) FSC curves (half-map FSC black, model vs. map FSC red). (**C**) Local resolution of the cryo-EM reconstruction. (**D**) Analysis of the cryo-EM map using the 3D FSC server (Tan et al., 2017) indicates isotropic resolution (sphericity = 0.9).

**Table S1. Cryo-EM data collection, 3D reconstruction, and refinement statistics.**

| **Dataset** | **CAK wild type** | **CAK-MAT1Δ219-THZ1** |
|---|---|---|
| Microscope | Talos Arctica | Talos Arctica |
| Stage type | Autoloader | Autoloader |
| Voltage (kV) | 200 | 200 |
| Detector | Gatan K3 | Gatan K3 |
| Acquisition mode | Super-resolution | Super-resolution |
| Physical pixel size (Å) | 0.72 | 0.72 |
| Defocus range (µm) | 0.3-2.5 | 0.5-2.5 |
| Electron exposure ($e^-/A^2$) | 69 | 69 |
| **Reconstruction** | **EMD-XXXX** | **EMD-XXXX** |
| Software | RELION 3.1 | RELION 3.1 |
| Particles picked | 8,982,448 | 4,884,787 |
| Particles final | 136,859 | 31,198 |
| Extraction box size (pixels) | 384 x 384 x 384 | 384 x 384 x 384 |
| Rescaled box size (pixels) | 216 x 216 x 216 | 192 x 192 x 192 |
| Final pixel size (Å) | 1.28 | 1.44 |
| Accuracy rotations (°) | 0.98 | 1.13 |
| Accuracy translations (Å) | 0.35 | 0.43 |
| Map resolution (Å) | 2.9 | 3.4 |
| Map resolution range | 2.85-3.25 | 3.25-3.9 |
| Map sharpening B-factor ($Å^2$) | -45 | -40 |
| **Coordinate refinement** | | |
| Software | PHENIX | PHENIX |
| Refinement algorithm | REAL SPACE | REAL SPACE |
| Clipped box size (pixels) | 144 | 128 |
| Resolution cutoff (Å) | 2.95 | 3.4 |
| $FSC_{model-vs-map}$=0.5 (Å) | 3.1 | 3.6 |
| **Model** | **PDB-XXXX** | **PDB-XXXX** |
| Number of residues | 646 | 645 |
| Protein | 644 | 644 |
| Ligand (ATPγS, THZ1, $Mg^{2+}$) | 2 | 1 |
| B-factors overall | 49.9 | 69.1 |
| Protein | 49.9 | 69.1 |
| Ligand (THZ1) | - | 81.0 |
| Ligand (ATPγS-$Mg^{2+}$) | 83.2 | - |
| R.M.S. deviations | | |
| Bond lengths (Å) | 0.006 | 0.005 |
| Bond angles (°) | 0.632 | 0.595 |
| **Validation** | | |
| Molprobity score | 1.9 | 2.2 |
| Molprobity clashscore | 3.6 | 5.7 |
| Rotamer outliers (%) | 3.9 | 5.0 |
| $C_β$ deviations (%) | 0 | 0 |
| Ramachandran plot | | |
| Favored (%) | 95.1 | 94.9 |
| Allowed (%) | 4.9 | 5.1 |
| Outliers (%) | 0.0 | 0.0 |

**Table S2. Components of the refined atomic models.**

| Protein | Chain | Length (aa) | Residues built | Ligand | Comments |
|---|---|---|---|---|---|
| | | | **CAK** | | |
| MAT1 | H | 309 | 244-308 | | |
| Cyclin H | I | 323 | 1-38, 42-284 | | |
| CDK7 | J | 346 | 10-45, 51-312 | ATPγS-$Mg^{2+}$, residues 400, 401 | S164-phosphate |
| | | | **CAK-THZ1** | | |
| MAT1 | H | 309 | 244-308 | | Residues 1-219 deleted |
| Cyclin H | I | 323 | 1-38, 42-284 | | |
| CDK7 | J | 346 | 10-45, 51-312 | THZ1, residue 401 | S164-phosphate |

**SI References**


Baumli S., Lolli G., Lowe E. D., Troiani S., Rusconi L., Bullock A. N., Debreczeni J. E., Knapp S., Johnson L. N. (2008) The structure of P-TEFb (CDK9/cyclin T1), its complex with flavopiridol and regulation by phosphorylation. *EMBO J* **27** (13)**:** 1907-1918

Jeffrey P. D., Russo A. A., Polyak K., Gibbs E., Hurwitz J., Massagué J., Pavletich N. P. (1995) Mechanism of CDK activation revealed by the structure of a cyclinA-CDK2 complex. *Nature* **376** (6538)**:** 313-320

Lolli G., Lowe E. D., Brown N. R., Johnson L. N. (2004) The crystal structure of human CDK7 and its protein recognition properties. *Structure* **12** (11)**:** 2067-2079

Robert X., Gouet P. (2014) Deciphering key features in protein structures with the new ENDscript server. *Nucleic Acids Res* **42** (Web Server issue)**:** W320-324

Russo A. A., Jeffrey P. D., Pavletich N. P. (1996) Structural basis of cyclin-dependent kinase activation by phosphorylation. *Nat Struct Biol* **3** (8)**:** 696-700

Schulze-Gahmen U., De Bondt H. L., Kim S. H. (1996) High-resolution crystal structures of human cyclin-dependent kinase 2 with and without ATP: bound waters and natural ligand as guides for inhibitor design. *J Med Chem* **39** (23)**:** 4540-4546

Sievers F., Wilm A., Dineen D., Gibson T. J., Karplus K., Li W., Lopez R., McWilliam H., Remmert M., Söding J., Thompson J. D., Higgins D. G. (2011) Fast, scalable generation of high-quality protein multiple sequence alignments using Clustal Omega. *Molecular Systems Biology* **7** (1)**:** 539

Tan Y. Z., Baldwin P. R., Davis J. H., Williamson J. R., Potter C. S., Carragher B., Lyumkis D. (2017) Addressing preferred specimen orientation in single-particle cryo-EM through tilting. *Nat Meth* **14** (8)**:** 793-796
